# Identification and targeting of a pan-genotypic influenza A virus RNA structure that mediates packaging and disease

**DOI:** 10.1101/2021.08.21.457170

**Authors:** Rachel J. Hagey, Menashe Elazar, Siqi Tian, Edward A. Pham, Wipapat Kladwang, Lily Ben-Avi, Khanh Nguyen, Anming Xiong, Meirav Rabinovich, Steven Schaffert, Talia Avisar, Benjamin Fram, Ping Liu, Purvesh Khatri, Jeffery K. Taubenberger, Rhiju Das, Jeffrey S. Glenn

## Abstract

Currently approved anti-influenza drugs target viral proteins, are subtype limited, and are challenged by rising antiviral resistance. To overcome these limitations, we sought to identify a conserved essential RNA secondary structure within the genomic RNA predicted to have greater constraints on mutation in response to therapeutics targeting this structure. Here, we identified and genetically validated an RNA stemloop structure we termed PSL2, which serves as a packaging signal for genome segment PB2 and is highly conserved across influenza A virus (IAV) isolates. RNA structural modeling rationalized known packaging-defective mutations and allowed for predictive mutagenesis tests. Disrupting and compensating mutations of PSL2’s structure give striking attenuation and restoration, respectively, of *in vitro* virus packaging and mortality in mice. Antisense Locked Nucleic Acid oligonucleotides (LNAs) designed against PSL2 dramatically inhibit IAV *in vitro* against viruses of different strains and subtypes, possess a high barrier to the development of antiviral resistance, and are equally effective against oseltamivir carboxylate-resistant virus. A single dose of LNA administered 3 days after, or 14 days before, a lethal IAV inoculum provides 100% survival. Moreover, such treatment led to the development of strong immunity to rechallenge with a ten-fold lethal inoculum. Together, these results have exciting implications for the development of a versatile novel class of antiviral therapeutics capable of prophylaxis, post-exposure treatment, and “just-in-time” universal vaccination against all IAV strains, including drug-resistant pandemics.

**One Sentence Summary:** Targeting a newly identified conserved RNA structure in the packaging signal region of influenza segment PB2 abrogates virus production *in vitro* and dramatically attenuates disease *in vivo*.

## Introduction

Influenza A virus (IAV) is a segmented RNA virus that causes major morbidity and mortality worldwide. Current antiviral therapies target viral proteins that frequently mutate, rendering many such therapies inadequate^1–4^. Despite a breadth of knowledge about the viral lifecycle, knowledge of the RNA secondary structure of the genome is limited. Recent research on other RNA viruses has revealed genomic RNA to be capable of playing many important roles in viral lifecycles beyond merely encoding amino acid sequences, suggesting that viral RNA structural elements could be promising therapeutic targets^5,6^. To the extent these RNA structural elements are both essential and highly conserved, these features could reduce the degree of freedom for mutations that are compatible with virus function. This in turn, could translate into a high barrier for resistance to therapeutics designed to disrupt these RNA structures. In IAV, genome packaging is one such critical juncture in which RNA structure might serve a central function.

The IAV genome consists of eight single-stranded negative-sense viral RNA (vRNA) segments that encode a minimum of 14 known viral proteins^7^. The vRNA, together with nucleoprotein (NP) and the heterotrimeric polymerase complex, comprised of PB2, PB1, and PA proteins, forms the complete viral ribonucleoprotein (vRNP)^8,9^. To be fully infectious, IAV virions must incorporate at least one of each segment’s vRNP^10,11^. Current paradigm supports a selective packaging method whereby the eight vRNPs are selected in a hierarchal, segment-dependent manner mediated by unique, segment-specific packaging signals present in the terminal and central coding regions of each segment that allow for discrimination between the vRNAs^9,12,13^. Each vRNP interacts with at least one other partner to form a supramolecular complex^14^ likely maintained by intersegment RNA-RNA and/or protein-RNA interactions hypothesized to guide the packaging process^10,15^. The mechanism mediating this selection and arrangement, however, is poorly understood. Curiously, packaging signals exist in regions of high nucleotide conservancy that strongly suppress synonymous codon usage^16–18^. Conservation of primary sequence beyond what is required for protein coding suggests the potential for maintenance of RNA structures possessing biological functionalities. Certain synonymous mutations within the polymerase gene, PB2, not only affect its own packaging, but also the incorporation of other segments^11,13,16^. We hypothesized that PB2’s dominant role in the packaging process might be facilitated by non-protein elements encoded by the PB2 RNA, including structured RNA elements. To test this hypothesis, we first solved the RNA secondary structure within PB2 that mediates packaging. We then genetically validated this structure’s critical role in the viral life cycle *in vitro* and IAV pathogenesis *in vivo*. Finally, we show proof-of-concept for a new class of antiviral therapeutics that can efficiently disrupt packaging and completely prevent and treat otherwise lethal disease *in vivo*, as well enable the development of strong functional immunity, with a high barrier to resistance.

## Results

### SHAPE-characterization of IAV segment PB2 packaging signal identifies conserved structure

To search for structured RNA domains, we first applied <u>s</u>elective 2′-<u>h</u>ydroxyl <u>a</u>cylation analyzed <u>b</u>y <u>p</u>rimer <u>e</u>xtension “SHAPE”^19^ and computational modeling to IAV segment PB2 genomic vRNA. *In vitro* transcribed full-length (-)-sense PB2 vRNA from strain A/Puerto Rico/8/1934 (H1N1) “PR8” was folded in solution^20^ and interrogated using an electrophilic SHAPE reagent that preferentially reacts with nucleotides existing in flexible, single-stranded states^19^ (**Fig. 1**). This analysis revealed that much of the 2341-nt vRNA is largely unstructured (**Supplementary Fig.1a**), consistent with other bioinformatics studies that found higher potential for RNA secondary structure conservation in the (+)-sense over the (-)-sense RNA for all segments, including PB2^17,21^. However, these previous studies did not analyze the terminal coding regions (TCR), and instead stopped 80 nucleotides short of the PB2 5′ TCR’s end. SHAPE-guided modeling suggested several areas in this terminal region that contain stable RNA secondary structures, most notably a stem-loop motif, named herein as Packaging Stem-Loop 2 (PSL2) (**Fig. 1a** and **Supplementary Fig.1b**, nucleotides 34-87). This region included a set of nucleotides that were previously implicated in segment PB2 packaging through mutational analysis via an unidentified mechanism (**Fig. 1a,b**, see circled nucleotides, and **Supplementary Table 1**)^13,16,18^. Supporting the hypothesis that these prior mutations act through disruption of PSL2 structure, SHAPE analysis of the mutants yielded different conformations that all abrogated the wild-type PSL2 structure **(Fig. 1c**, and **Supplementary Fig. 2**). The 60-nucleotide region encompassing PSL2 displays near 100% sequence conservation at the single nucleotide level between representative seasonal as well as pandemic strains of different subtypes and species origins (**Fig. 1d, Supplementary Fig. 1c**). Further analysis revealed that this high degree of conservation extends to all known IAV isolates available in public databases (**Supplementary Fig**.**1d**), suggesting the existence of a strict biologic requirement to maintain an intact PSL2 structure. To exclude the possibility that differing downstream sequences within PB2 vRNA could alter the secondary structure of PSL2, we explored PSL2’s structural conservancy by performing SHAPE on full-length wild-type PB2 vRNAs from a variety of IAV strains and subtypes, including the highly pathogenic avian H5N1 and pandemic 1918 H1N1 strains. Despite the presence of two diverging nucleotides within the stem-loop and significant divergence in flanking sequences, the PSL2 stem-loop structure was recovered in SHAPE-guided modeling of PB2 RNA across these diverse species and subtypes (**Fig. 1e**).

**Figure 1.**
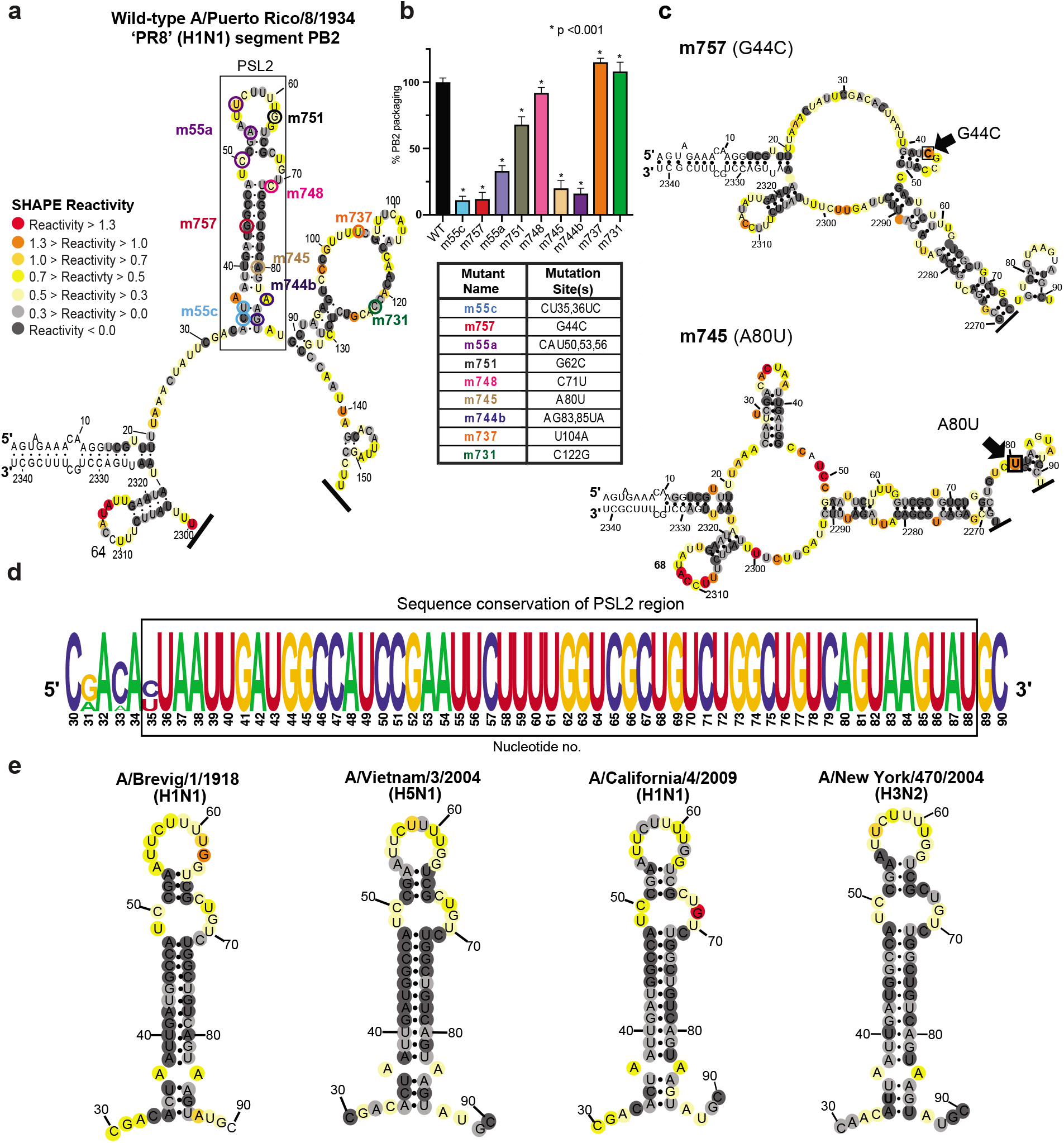
SHAPE-determined RNA secondary structures of wild-type PB2 and packaging mutant vRNAs. SHAPE-chemical mapping performed on full-length (-)-sense PB2 vRNAs. Colors denote SHAPE reactivity, which is proportional to the probability that a nucleotide is single-stranded. All structures are truncated to highlight the 5′ termini sequence structure. (**a**) SHAPE-predicted wild-type PB2 RNA secondary structure from strain A/Puerto Rico/8/1934 “PR8” (H1N1). Color-coded circles correspond to nucleotides sites where synonymous mutations were reported to affect PB2 packaging^13,16^. (**b**) Packaging efficiency of synonymous mutants in (a), determined by qPCR. Results representative of two independent experiments with biological replicates, each performed in triplicate. Statistical analysis was performed using one-way ANOVA with Dunnett’s multiple comparisons test against the WT mean by GraphPad Prism software. Error bars represents ± standard deviation (s.d.). Box below indicates mutant name and corresponding nucleotide change. Nucleotide numbering shown in the genomic (-)-sense orientation. (**c**) SHAPE-determined structures of PB2 packaging-defective mutant vRNAs, m757 (G44C) and m745 (A80U) indicating loss of PSL2’s RNA secondary structure. Black arrowheads and boxed nucleotides denote site(s) of synonymous mutation. (**d**) Web-logo representation of the PSL2 region conservation across IAV strains and diverse influenza A viral subtypes (weblogo.berkeley.edu). The overall height represents sequence conservation at that nucleotide position, while the height of symbols within each position indicates the relative frequency of each nucleotide at that site. Black box denotes PSL2 region. Sequences included in the alignment: pandemic A/Brevig Mission/1/1918 (H1N1), pandemic A/California/04/2009 (H1N1), seasonal human A/New York/470/2004 (H3N2), A/Puerto Rico/8/1934 (H1N1), high pathogenic avian A/Vietnam/03/2004 (H5N1), avian A/mallard/Maryland/14OS1154/2014 (H6N1), pandemic A/Hong Kong/8/1968 (H3N2), and seasonal human A/New York/312/2001 (H1N1) (*see* Supplementary Fig. 1c). RNA nucleotides are numbered in (-)-sense orientation. (**e)** SHAPE-determined structures of wild-type PB2 vRNA from pandemic and highly pathogenic strains, including different subtypes to modern human strains: 1918 pandemic (A/Brevig Mission/1/1918 (H1N1)), highly-pathogenic avian (A/Vietnam/1203/2004 (H5N1)), 2009 pandemic ‘swine’ (A/California/04/2009 (H1N1)), and Fujian-like human seasonal virus, A/New York/470/2004 (H3N2).

### Mutate-and-Map strategy validates PSL2 structure and predicts novel packaging mutants

To further test the SHAPE analysis of the PSL2 RNA structure and to uncover additional informative mutations needed for *in vivo* tests, we applied multidimensional chemical mapping methods to the PSL2 segment. Mutate-and-Map (M^2^) analysis couples systematic mutagenesis with high-throughput chemical mapping to produce accurate base-pair inferences and interactions of RNA domains^22^. By sequentially mutating RNA one nucleotide at a time with its Watson-Crick complement and measuring the impact this mutagenesis has on chemical reactivity, pair-wise correlations between close and distant residues can be established. First, M^2^ measurements confirmed disruption of the chemical reactivity pattern upon systematic mutation of each PSL2 stem residue, including changes at nucleotides previously identified to be relevant for PB2 packaging (**Fig. 2**, *see* noted fields)^13,16^. Automated computational analysis based on these M^2^ data recovered the SHAPE-guided PSL2 structure with high confidence (**Fig. 2, Supplementary Figs. 2** and **3**), further validating our structural model. Second, as predictive tests, we designed compensatory mutations to restore base pairings—albeit not the native sequence—in the wild-type stem-loop structure that were disrupted by the initial packaging-defective mutations (**Supplementary Fig. 4**). These mutation-rescue variants indeed restored the PSL2 SHAPE pattern, providing *in vitro* validation of the modeled structure at base-pair resolution and suggesting sequence variants to test the role of PSL2 structure *in vivo*.

**Figure 2.**
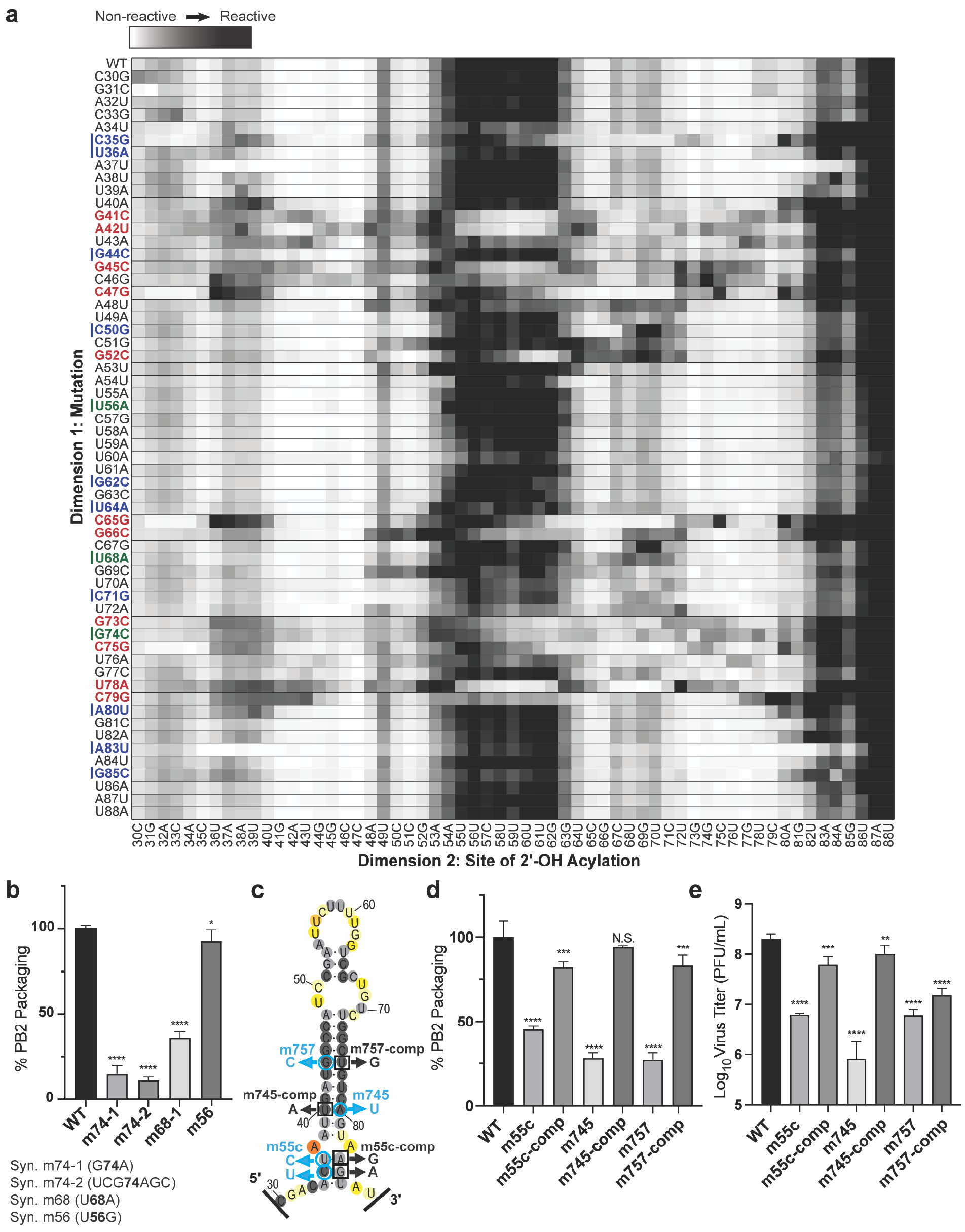
2-Dimensional Mutate-and-Map (M^2^) analysis and empiric validation of PSL2 motif. (**a**) Systematic single nucleotide mutation and mapping of resulting chemical accessibility reveals interactions in the three-dimensional structure of the RNA. Chemical accessibilities, plotted in grayscale (black, highest SHAPE reactivity), across 59 single mutations at single-nucleotide resolutions of PSL2 element from PR8 strain segment PB2. Reactivity peaks (left to right) correspond to nucleotides from the 5′ to 3′ end of the PB2 RNA. Nucleotides corresponding to known packaging mutation sites (ref. *13*) are indicated on left in blue. Red bolded mutations denote prominent packaging-defective mutant sites predicted by M^2^ analysis. Green bolded mutations indicate synonymous mutant sites analyzed in (b). (**b**) Packaging efficiencies of M^2^-identified synonymous mutants read out by qPCR. Packaging efficiency represents the percentage of mutant PB2 vRNA packaging relative to parental wild-type PB2. Results from two independent experiments in biological duplicate, performed in technical triplicate (n=4). Statistical analysis performed by ordinary one-way ANOVA using Dunnett’s multiple comparisons test against WT computed in GraphPad Prism software. (**c**) Previously described synonymous mutants (m757, m745, m55c) are mapped onto PSL2 structure. Compensatory, non-synonymous mutations m55c-comp, m745-comp, and m757-comp were designed at sites predicted to restore wild-type PSL2 structure based on SHAPE and mutate-and-map chemical analyses. Black boxed nucleotides denote site of compensatory mutation. (-)-sense vRNA orientation is shown. (**d**) Packaging efficiencies of packaging-defective and compensatory PB2 mutant viruses. For compensatory mutations where a non-synonymous change was required, a wild-type PB2 protein expression plasmid was co-transfected during virus rescue. Values given as percentage of PB2 vRNA packaging in comparison to wild-type parental PR8 virus. Results from three independent experiments, assays performed in triplicate. (**e**) Virus titer determined by plaque assay. Results in PFU / mL, plaque assays performed in triplicate. All error bars represent ± s.d. Statistical analyses in (d-e) performed as stated in (b) above. * p < 0.05 ** p <0.01; *** p < 0.001; **** p < 0.0001.

To test whether the PSL2 stem-loop structure observed in solution was relevant to virus packaging in the cellular milieu, the same nine synonymous mutations reported by Gog et al. (2007) and Marsh et al. (2008) (**Fig. 1a,b** and **Supplementary Table 1**) as well as four new synonymous mutations characterized by M^2^ analysis (**Fig. 2a,b**) were cloned into pDZ plasmids containing the PR8 PB2 gene^13,16,18^. The packaging efficiencies of the nine previously known mutants, now in the PR8 background, were comparable to those originally described in the WSN33 virus^13^ (**Fig. 1b** and **Supplementary Table 1).** Of these, mutants m55c, m757, m745, and m744b, were predicted to show the most significant impairment based on their location within PSL2’s stem regions (**Fig. 1a-c** and **Supplementary Fig. 2**). In contrast, published mutations that have no effect on PB2 packaging (e.g. m731) mapped to the unstructured apical loop or fell outside of PSL2 and did not alter its structural integrity (**Supplementary Fig. 5a**), while mutations with minor effects to virus packaging showed only minor alterations to the structure (**Supplementary Fig. 5b**) ^13^. The three novel synonymous mutants (m74-1, m74-2, and m68) identified by M^2^-analysis as having a significant effect on *in vitro* PSL2 structure (**Fig. 2a**, *see* green-marked nucleotides) showed significant loss in PB2 packaging, whereas mutation sites that resulted in negligible change in SHAPE reactivity compared to wild-type PSL2 (e.g., m56), gave wild-type-like packaging efficiency levels (**Fig. 2b**). The strong correlation of structure disruption with *in cellulo* packaging efficiency observed across these mutants supports a role of PSL2 structure in virus packaging.

### Compensatory mutation pairs restore PSL2 structure, rescue packaging in vitro and in vivo disease

To investigate the functional role of PSL2 in IAV genome packaging, compensatory mutations designed to restore the wild-type stemloop structure destroyed by the packaging-defective mutations (**Fig. 2c, Supplementary Fig. 4**) were cloned into PR8 pDZ plasmids to generate mutant rescue viruses. The compensatory mutations rescued not only the virus packaging for segment PB2 (**Fig. 2d**), but also other segments previously reported to be affected by the deleterious mutations, consistent with the proposed hierarchal role of PB2 in IAV packaging (**Supplementary Fig. 6a**)^11–13^. In addition to recovering PB2 packaging, the compensatory mutations gave complete or near-complete rescue of the virus titer loss caused by the defective mutations (**Fig. 2e** and **Supplementary Fig. 6b**). Some non-synonymous compensatory mutations were able to restore PB2 packaging better than others (m745-comp and m55c-comp, compared to m757-comp). This possibly reflects incomplete restoration of PB2 protein function through exogenous addition (**Fig. 2d, e** and **Supplementary Fig. S6**) since for non-synonymous mutations, we also expressed WT PB2 protein to mitigate the possibility of any impairment in PB2 protein function.

The most incisive test of PSL2 structure came from packaging experiments that did not require supplemental wild-type PB2 protein addition. Computational enumeration and multidimensional mutation-rescue^23^ (M^2^R) experiments were performed in order to identify additional successful PSL2-defective and compensatory mutant pairs (**Fig. 3a, b** and **Supplementary Fig. 7**). Successful mutation-rescue was defined when each single mutation alone resulted in disrupting the SHAPE-mapped wild-type PSL2 structure, while the double compensatory mutations recovered wild-type PSL2 structure (**Fig. 3a, b** *see* boxed electropherograms). Although most discovered successful partners required non-synonymous changes, we discovered a single mutation-rescue pair of substitutions that were both synonymous, obviating wild-type PB2 protein addition (**Fig. 3b, c** and **Supplementary Figs. 7, 8** and **9**). Making each mutation alone (m52 and m65) resulted in severe packaging defects and virus titer loss exceeding 4 log_10_—an extreme impairment beyond what has been previously reported^13,16^ (i.e. 1-2 log_10_) for packaging-defective viruses (**Figs. 3d, e** and **Supplementary Figs. 6** and **8**). When introduced together into a doubly mutated m52/65-compensatory strain that restored PSL2 structure, albeit with an altered sequence, the compensatory mutations restored both packaging efficiency and virus titer to wild-type levels. These data provided unambiguous evidence for the PSL2 52-65 RNA base pair in influenza A packaging. In order to ensure that any loss or subsequent rescue of virus packaging was not due to defects in replication or translation caused by these synonymous mutations, each of the PB2 mutation-rescue mutants were tested in a transfection-based replicon assay to assess both the ability of the generated PB2 protein mutants to function as part of the polymerase complex, as well as to test the ability of the PB2 mutant vRNA to be replicated. All mutant PB2 proteins and vRNAs were produced at comparable wild-type levels (**Supplementary Figs. 10** and **11**).

**Figure 3.**
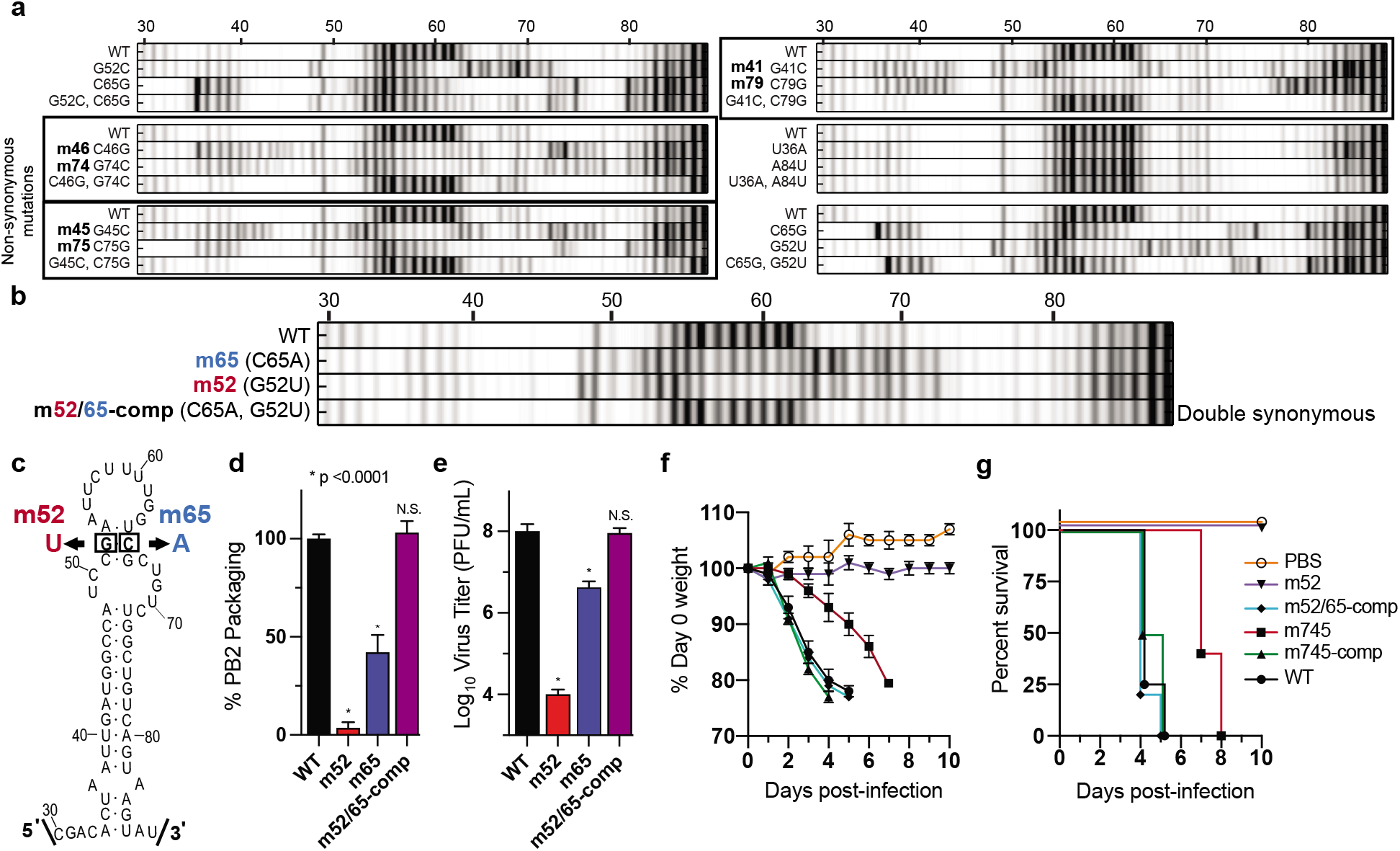
Mutate-Map-Rescue analysis reveals novel PB2 packaging-defective and compensatory mutant partners. (**a**) Electropherograms from systematic single nucleotide mutation SHAPE chemical mapping with rescue (Mutate-Map-Rescue)^21^ analysis of individual and compensatory double mutations to test base-pairings from 1D-data-guided models and to identify predicted successful synonymous PSL2-defective and compensatory mutant pairs. Chemical accessibilities, plotted in grayscale (black = highest SHAPE reactivity), across 59 single mutations at single-nucleotide resolutions of PSL2 element from PR8 strain segment PB2. Reactivity peaks (left to right) correspond to nucleotides from the 5′ to 3′ end of the PB2 RNA. *See* Supplementary Fig. 7 for complete set of Mutate-Rescue pairs. (**b**) Electropherogram of successful double synonymous mutant pair determined by Mutate-Map-Rescue analysis. (**c**) Mutational design of single mutants m52 (G52U) and m65 (C65A), and the double m52/65-comp rescue pair on the PSL2 structure. (**d**) Packaging efficiency of the synonymous single and double mutant Mutate-Rescue pair. Values given as a percentage of PB2 vRNA packaging relative to WT parental PR8 virus. Results represent two independent experiments with biological replicates and performed in technical triplicate (n=4) except for the m65 mutant, which were performed in biological triplicate (n=6). (**e**) Viral titer of the supernatants collected in (d) in PFU / mL, plaque assay results in triplicate. Statistical analysis performed in (d-e) by ordinary one-way ANOVA using Dunnett’s multiple comparisons test against WT using GraphPad Prism software. * p <0.0001, N.S. = not significant. (**f-g**) Percent Day 0 weight and survival of mice infected with single PSL2-disrupting, and compensatory PSL2-restoring double-mutant viruses. Six to eight weeks old BALB/c female mice (n=6 mice / group) were intranasally infected with PR8 wild-type (WT) virus, packaging-defective single mutant viruses, m52 and m745, compensatory double mutant viruses, m52/65-comp and m745-comp, or PBS control. (**f**) Percent Day 0 weight. (**g**) Kaplan-Meier survival plot of the individual cohorts depicted in (f). All error bars represent ± s.d.

To test the relevance of the PSL2 structure in an *in vivo* model, 6-8 week old BALB/c mice were intranasally instilled with either wild-type or mutant PR8 viruses harboring point mutations predicted to disrupt or restore PSL2 structure. Mice infected with the PSL2-disrupting mutations—m745 mutant strain (20% packaging efficiency) or the severely packaging-defective single mutant virus, m52 (<4% packaging efficiency)—showed reduced or no clinical signs of illness, respectively, either in weight loss or survival as compared to the PBS control (**Fig. 3f, g**). Remarkably, inclusion of compensatory mutations that restore PSL2 structure rescued virus pathogenicity: animals infected with m52/65-comp and m745-comp, displayed comparable mortality profiles as mice infected with wild-type PR8 (**Fig. 3f, g**). To the best of our knowledge, these are the first data indicating that packaging-defective viruses are attenuated *in vivo* and a genomic IAV RNA secondary structure mediates influenza disease progression. Given the strong evolutionary conservation of the predicted PSL2 structure across different IAV subtypes, strains, and host species isolates (**Fig. 1d, e** and **Supplementary Fig. 1c, d**), we postulated that therapeutics directed against this structure could possess broad-spectrum antiviral activity against all IAV subtypes and strains.

### Therapeutic design and antisense targeting of PSL2 structure inhibits IAV infection in vitro

To explore the therapeutic potential of targeting PSL2-mediated virus packaging, nine antisense oligonucleotides (ASO) with modified locked nucleic acid (LNA) bases containing phosphorothioate internucleoside linkages^24^ were designed against PSL2 residues in a manner predicted to disrupt various aspects of the overall RNA secondary structure of the motif and their effect on inhibition of virus production was determined (**Fig. 4a**). Two of the designed LNAs, LNA8 and LNA9, are identical in sequence to LNA6 and LNA7, respectively, but possess 6-7 unmodified (non-locked) DNA nucleotide “gapmers” optimized for RNase-H activation that can degrade RNA in RNA-DNA hybrids^25^. First, to assess the impact that LNA binding has on PSL2 RNA secondary structure, toeprinting and SHAPE chemical mapping were performed on PB2 vRNA in the presence of the LNAs. Sequences encoded in LNAs 6-9, corresponding to binding sites on the right 3’ side of the stemloop structure (**Fig. 4a**), exhibited the greatest ability to bind and disrupt the wild-type PSL2 structure (**Supplementary Fig. 12**).

**Figure 4.**
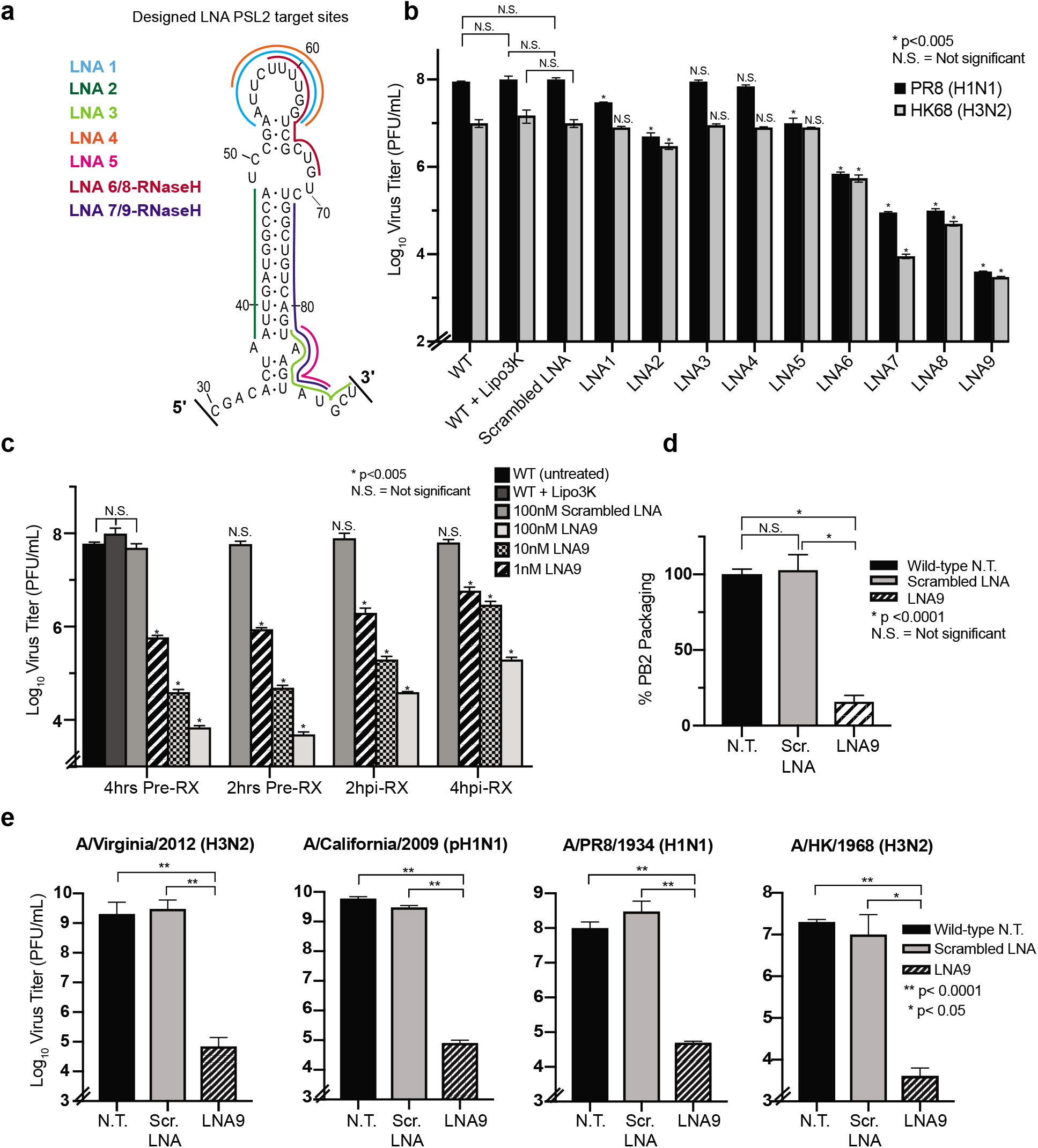
Locked Nucleic Acids targeting PSL2 RNA structure display potent antiviral activity *in vitro*. (**a**) Location of complementary Locked Nucleic Acids (LNAs) designed against different regions of the PSL2 structure. (**b**) LNA antiviral screen: MDCK cells were pretreated with 100 nM of each designated LNA or a scrambled mismatch LNA for 4 hours prior to infection with PR8 (H1N1) virus or A/Hong Kong/8/68 (H3N2) virus (0.01 MOI). 48 hours post-infection (hpi), supernatant was collected, and viral titers determined by plaque assay. Assays performed in biological triplicate with three technical replicates (n=9). * Statistical significance shown between non-treated WT + Lipo and LNA-treated samples, unless otherwise indicated. (**c**) Time course of pre-treatment (Pre-RX) versus post-infection treatment with LNA9 at titrating concentrations (100 nM, 10 nM, 1 nM). WT + Lipo3k = non-treated infection with Lipofectamine 3000 control. Pre-RX: MDCK cells were treated with LNA9 or Scrambled LNA either 2 or 4 hours prior to infection with PR8 virus at an MOI of 0.01. Post-infection treatment: MDCK cells were infected with PR8 virus at an MOI of 0.01 and treated with LNA9 or Scrambled LNA at either 2 or 4 hpi. Supernatant was collected 48 hpi, and viral titers determined by plaque assay in biological and technical triplicate (n=9). * Significance determined relative to WT PR8-Lipo. (**d**) Packaging efficiency of PB2 vRNA from PR8 viruses treated with 100 nM LNA9 or scrambled LNA control. Values given as a percentage of PB2 vRNA packaging in comparison to non-treated wild-type PR8 virus, readout by qPCR. Results from two biological replicates, assays performed in technical triplicates (n=6). (**e**) LNA9 antiviral efficacy against multiple influenza A strains: A/Virginia/2012 (H3N2), A/California/2009 (pH1N1), A/PR8/1934 (H1N1), and A/Hong Kong/1968 (H3N2). MDCK cells were pretreated with 100 nM of LNA9 or Scramble 12 hours prior to infection with the indicated viruses at an MOI of 0.01. Plaque assays performed in biological and technical triplicate (n=9). All error bars represent ± s.d.

To test the antiviral potential of LNA-mediated targeting of PSL2 across different IAV subtypes, MDCK cells were pre-treated with 100 nM of each LNA for 4 hours prior to infection with either the wild-type PR8 (H1N1) virus or the tissue culture-adapted A/Hong Kong/8/1968 (HK68) (H3N2) virus. Forty-eight hours post-infection, the supernatants were collected, and virus production was measured by plaque assays (**Fig. 4b**). As predicted by our mutational and LNA chemical mapping experiments (**Fig. 2, Supplementary Fig. 12**), LNAs directed against only the top loop of PSL2 (LNA1, LNA4), and LNAs solely targeting the 3′ base of PSL2 (e.g., LNAs 3 and 5) had minimal effect on viral titer. In contrast, nucleotide coverage of both the top loop and middle bulge by LNA6 resulted in greater than 2 log_10_ titer deficits for PR8 (**Fig. 4a, b**). LNA8, the RNase-H activated copy of LNA6, produced even greater antiviral activity against both viruses of up to 3 logs_10_. Most strikingly, LNA9, the RNase-H activated copy of LNA7, possessed the strongest antiviral capacity, dropping virus production by over 4 logs_10_ and 3 logs_10_ against PR8 and HK68, respectively. While LNA9 could be clearly visualized in cells harboring vRNPs (**Supplementary Fig. 13)**, no off-target effect of LNA9 on steady state levels of viral protein, vRNA, cRNA, or cellular toxicity after 24 hours was observed (**Supplementary Figs. 14, 15,** and **16**). The potent antiviral activity of these LNAs corroborate the results from the LNA chemical mapping experiment, indicating that other compounds that can similarly disrupt the PSL2 structure are likely to possess antiviral activity.

Having identified a potent candidate LNA, we next investigated the treatment time-course and concentration parameters of LNA9’s antiviral activity. MDCK cells were treated with varying concentrations of LNA9 at either 2 or 4 hours pre-infection or, alternatively, 2 or 4 hours post-infection with wild-type PR8 virus. Cells pre-treated with the LNA had the most potent antiviral response (greater than 4 logs_10_) and displayed strong virus inhibition (greater than 2 logs_10_) even at the lowest tested concentration (1 nM) (**Fig. 4c**). There was a trend towards decreasing antiviral activity as the time post-infection treatment increased, but even at the latest tested time point of addition, greater than 2 logs_10_ suppression of viral titer was achieved. Similar efficacy was seen in the presence of high MOI (**Supplementary Fig. 17**). To probe whether LNA-mediated virus inhibition resulted in loss of PB2 packaging, cells were treated with 100 nM of LNA9 or scrambled LNA and then infected with PR8 virus. Isolated vRNA from packaged virions were then analyzed for virus packaging by qPCR. Similar to the mutational studies, LNA9 treatment resulted in a dramatic loss of PB2 packaging compared to controls (**Fig. 4d**). To test the hypothesis that therapeutic targeting of the PSL2 motif would possess broad-spectrum antiviral activity, cells were pretreated with 100 nM of LNA9 or scrambled LNA prior to infection with more clinically relevant human strains, including the 2009 pandemic “swine” (pH1N1) virus. Both the modern H3N2 and pH1N1 viruses were highly sensitive to LNA9 treatment, showing inhibition at levels greater than or equal to results seen against PR8 or HK68 (**Fig. 4e**).

### In vitro selection of IAV variants under escalating drug pressure

Our designed LNAs are directed against a highly conserved genomic RNA target (**Fig. 1c, Supplementary Fig. 1c, d**) that appears to have strong biologic constraints against viable viral mutants. We hypothesized that this additional level of mutational constraint would provide a higher barrier to the development of resistance for therapeutics directed against PSL2. To test this hypothesis, we determined the susceptibility of wild-type PR8 virus to LNA9 under conditions designed to promote the development of resistance over serial virus passaging (**Fig. 5a**). In parallel, we performed analogous experiments using the neuraminidase inhibitor (NAI) oseltamivir carboxylate (OSLT, Tamiflu™). OSLT had a starting EC_50_ of 4.1 nM against PR8 at passage 1 of drug treatment. After seven virus passages in the presence of escalating drug concentrations, the EC_50_ of OSLT leapt to 100 µM—a greater than 20,000x fold increase (**Fig. 5b**). In comparison, after 10 passages of virus in the presence of LNA9, the EC_50_ held in the picomolar range of 16 to 22 pM. To date, we have yet to be able to select for viral mutations capable of generating resistance to LNA9. To test if LNA9 is efficacious against drug-resistant viruses, a drug-resistant mutant of A/WSN/33 (H1N1) virus was generated using a reverse genetic virus rescue system that mutated the NA gene to contain the H275Y mutation (N1 numbering system), known to confer resistance to NAIs. Against this NAI-resistant virus, LNA9 maintained the same potency and efficacy it exerted against wild-type, while high-level resistance to OSLT was confirmed, with an EC_50_ of 53 μM (**Fig. 5 c-e**). This result extends the apparent therapeutic capabilities of LNA9 and provides strong support for potential therapeutic treatment of NAI-resistant viruses with PSL2-targeting LNAs.

**Figure 5.**
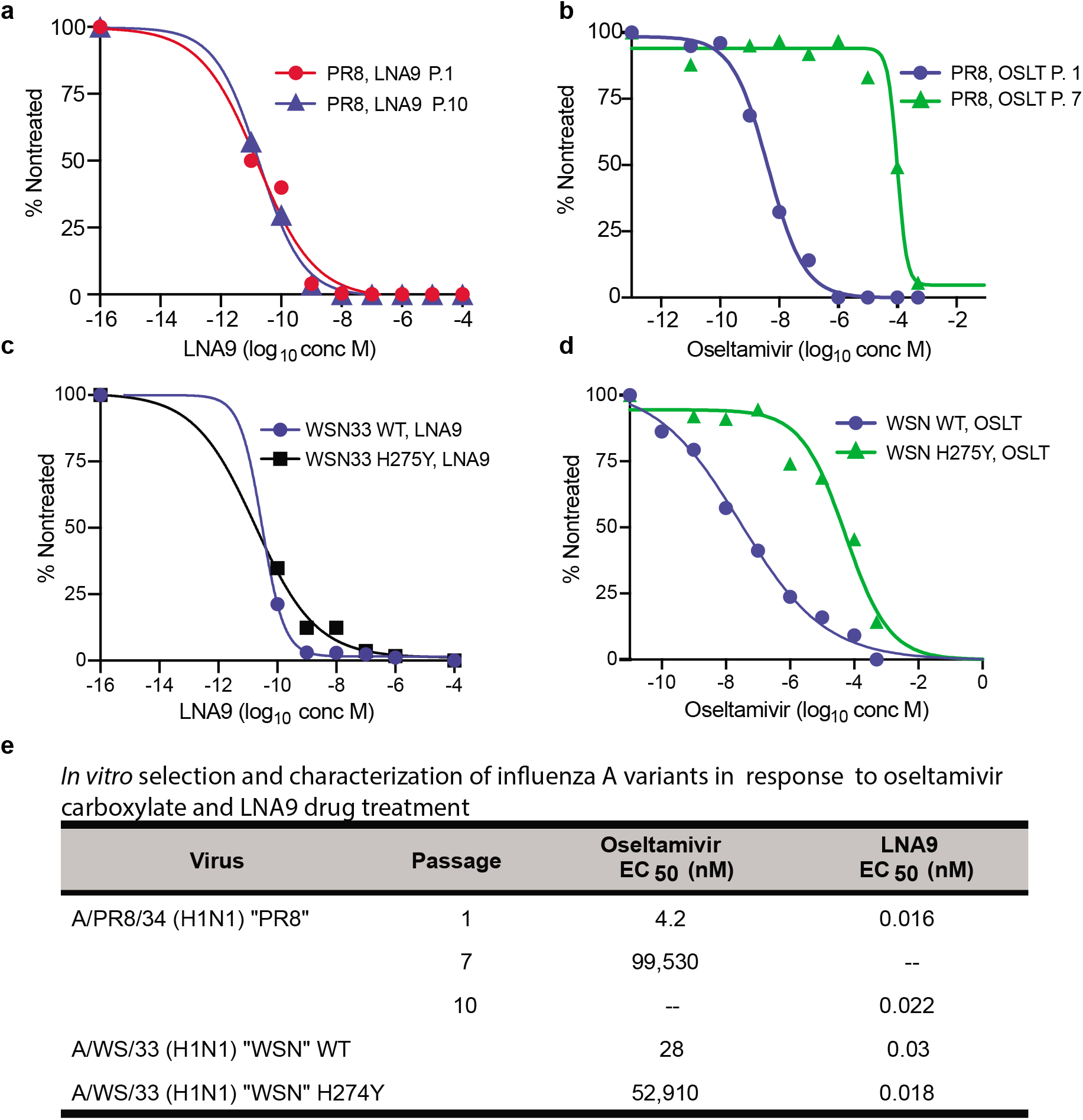
*In vitro* selection and sensitivity of passaged virus in response to drug treatment. (**a,b**) Wild-type PR8 virus was serially passaged in the presence of either LNA9 (a) or oseltamivir carboxylate (b) over time with escalating concentrations of drug. Viral supernatant from each passage was collected and titered by plaque assay. (**a**) LNA9: MDCK cells were pretreated with varying concentrations of LNA9 12 hours prior to infection at 0.01 MOI of passage 1 (P.1) or passage 10 (P.10) LNA9-treated PR8 virus. After 48 hours, viral supernatant was collected and titered for each drug dilution. Results expressed as a percentage of nontreated virus titer. The drug concentration that caused 50% decrease in the percent of PFU titer in comparison to untreated controls was defined as the EC_50_. (**b**) Oseltamivir: Confluent MDCK cells were infected with 100 PFU of passage 1 (P.1) or passage 7 (P.7) OSLT-treated PR8 virus and drug sensitivity was determined by plaque reduction assay. The number of viral plaques with each drug concentration was counted and plaque number was normalized against the nontreated control to determine the EC_50_. (**c,d**) *In vitro* sensitivity of wild-type WSN33 (H1N1) virus and NAI-resistant WSN H275Y NA mutant virus to LNA9 (c) or oseltamivir carboxylate (d). (**e**) Summary table of EC_50_ values from graphs (a-d).

### PSL2-targeted LNAs protect mice from lethal IAV infections

As a proof-of-concept to assess the *in vivo* efficacy of prophylactic LNA treatment against PSL2, BALB/c mice were intranasally (I.N.) administered a single 20 μg dose of LNA9 or scrambled LNA one day, or three days, prior to infection with a lethal dose of PR8 virus. The untreated control mice experienced dramatic weight loss and were humanely sacrificed by days 5 and 6. In contrast, a single dose of LNA9 was completely protective when administered one day, and even three days, prior to viral infection, and showed significantly reduced virus titer in the lungs (> 2.5 log_10_ virus reduction) compared to the scrambled control at 72 hrs post-infection (**Fig. 6a, Supplementary Fig. 18**). In addition to the well-known benefits of LNA antisense gapmer technology (e.g., high target binding affinity, RNase-H cleavage activity, and high nuclease resistance conferred by the thiolated phosphate backbone), LNA ASOs have also been reported to show dramatic, long-lasting effects (even >1 month) after the last administered dose in a variety of disease models^26–29^. We hypothesized that PSL2-targeted LNAs might similarly possess long-term prophylactic effects.

**Figure 6.**
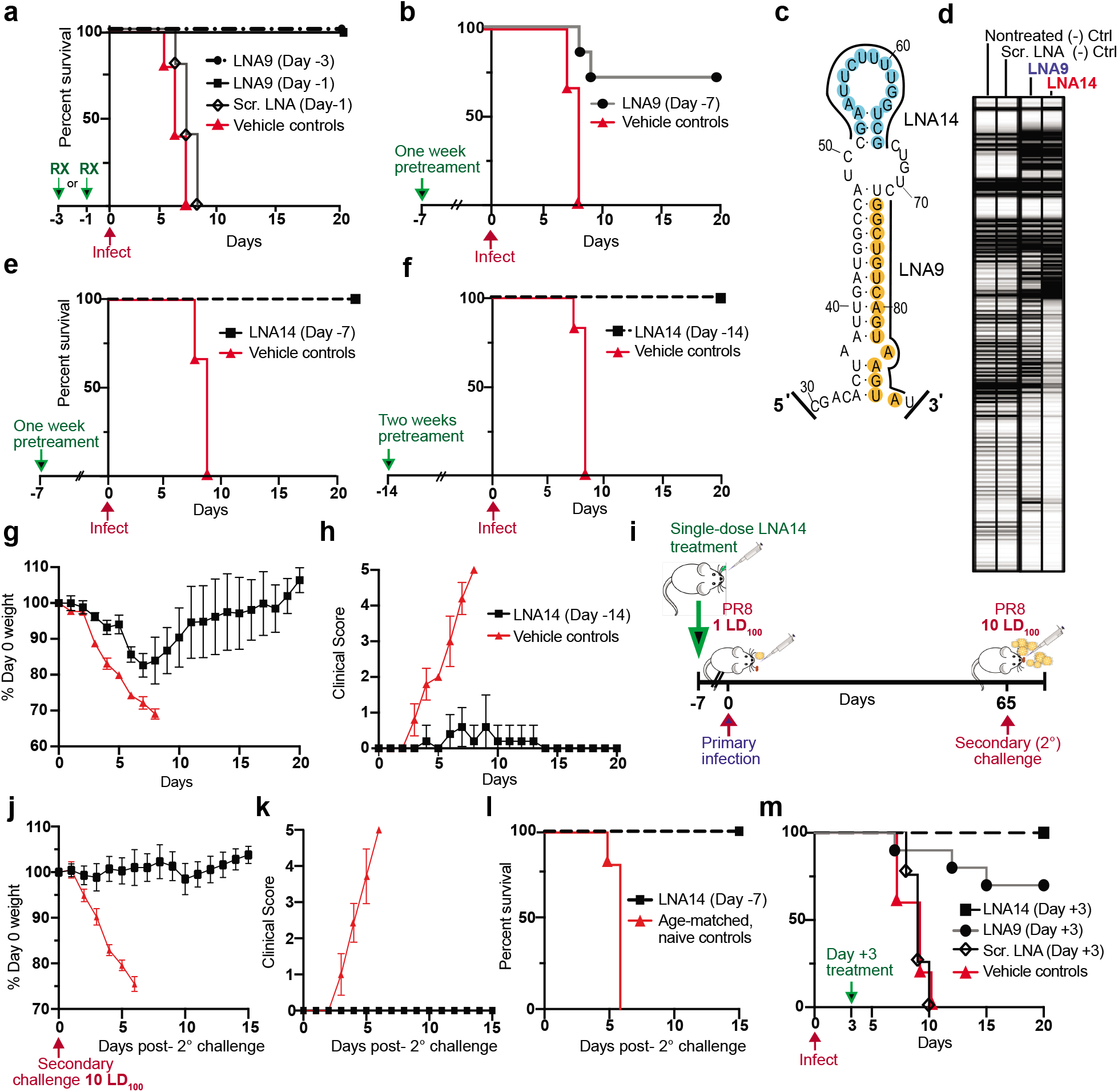
PSL2-targeted LNAs demonstrate potent antiviral activity *in vivo*. (**a-b**) Effect of intranasal LNA prophylactic treatment on survival of virus-infected mice. Kaplan-Meier survival plots. Mice (n=7 mice/group) were intranasally administered a single dose of LNA9, Scrambled LNA, or Vehicle (mock-treated) followed by a lethal inoculum of wild-type PR8 virus (**a**) dosed with 20 μg LNA 3 days before infection (Day −3) or 1 day before infection (Day −1); (**b**) one week pretreatment with a single 30 μg dose of LNA9 or vehicle control. (**c**) Target sites of LNA9 and newly designed, LNA14, mapped to the PSL2 structure. (**d**) Electrophoretic profile of SHAPE analysis performed on non-treated, Scrambled LNA, LNA9, and LNA14 at 100 nM concentration in the presence of PR8 PB2 vRNA. Labeling with 1M7 SHAPE reagent shown. (**e**) Kaplan-Meier survival plot of mice (n=7 mice/group) intranasally pretreated with a single dose of 30 μg LNA14 or vehicle control one week (Day −7) before lethal PR8 infection. (**f-h**) A single dose of 40 μg LNA14 or vehicle was I.N. administered two weeks (Day −14) before PR8 virus infection. (**f**) Kaplan-Meier survival plot. (**g**) Percent day 0 weight of mice. (**h**) Clinical score. (**i-l**) Mice (n=7) were given a single 40 μg intranasal dose of LNA14 one week prior to a primary lethal infection at 1 LD_100_ of PR8 virus (e). Sixty-five days post-initial infection, surviving mice from (e) along with age-matched naïve controls (n=7/group) were challenged a second time at 10 LD_100_. (**i**) Challenge study timeline. (**j**) Percent day 0 weight of mice. (**k**) Clinical score. (**l**) Kaplan-Meier survival curve. (**m**) Mice (n=10/group) were infected with a lethal dose of PR8 wild-type virus. Three days post-infection, mice were given a single dose of 40 μg of LNA14, LNA9, Scrambled LNA, or vehicle control by I.V. injection. Kaplan-Meier survival plot.

To test this hypothesis, we administered a single, increased dose of LNA9 (30 μg) one week (Day −7) prior to infection with a lethal dose of IAV (**Fig. 6b**). While 100% of untreated mice succumbed to the infection, 70% of the LNA9 one-week pretreatment group were protected from lethality. To determine if our therapeutics can be further optimized for improved efficacy, we fine-tuned the delivery formulation as well as reconsidered the LNA target region. While LNA9 targets nucleotides in the lower 3’ stem of PSL2, our mutational analyses suggest the importance of the 52-65 nucleotide pair in the upper stem (**Fig. 4**). In fact, none of LNAs1-9 fully covered this region, so we hypothesized that additional LNA design against these nucleotides might enhance efficacy (**Fig. 4a**). The newly designed LNA14 fully covers the top stemloop of PSL2, and upon SHAPE analysis comparing LNA activity against PSL2, we found LNA14 to be an even more potent disruptor of PSL2 structure than LNA9 (**Fig. 6c, d**). In biological confirmation of this, mice treated with LNA14 were fully protected when given a single dose one week prior to infection with a lethal dose of IAV (**Fig. 6e**). Pushing the prophylactic window even further, we again increased the LNA dosage (40 μg) and administered a single LNA14 treatment to mice two-weeks (Day −14) prior to IAV infection. While the mock-treated mice were sacrificed between days 7 and 8 due to severe disease, the entire LNA14-treated cohort survived, showing significantly lower clinical scores indicative of minor-to-undetectable disease symptoms and reduced weight loss compared to controls (**Fig. 6f-h, Supplementary Table 2**).

Anti-PSL2 LNA treatment protects mice from lethal infection in part by attenuating the virus and reducing disease progression; however, because mice do display symptoms, albeit minor, we hypothesized that the resulting highly attenuated infection might be sufficient to enable mice receiving prophylactic LNA treatment to develop an effective immunization against a secondary infection through production of long-term immunity. To test this hypothesis, mice (n=7) from the one-week LNA14 pretreatment surviving cohort (**Fig. 6e**) were challenged alongside age-matched, naïve controls, sixty-five days post-primary infection with ten times the mouse lethal dose (10 LD_100_) of IAV (**Fig. 6i**). The resulting secondary challenge had no effect on weight, clinical score, or survival of mice from the LNA14 pretreatment group (a total of 72 days since treatment, 65 days since primary infection), while the age-matched controls presented with rapid disease and were humanely sacrificed by Day 6 post-challenge infection (**Fig. 6j-l**).

After demonstrating PSL2-targeted LNA efficacy in prophylactic models, we next sought to establish its potential as a post-infection therapeutic. Due to the rapid onset of symptoms and illness in IAV infections, FDA-approved IAV therapeutics are most challenged when administered after 48 hrs of disease onset^30^. To assess the ability of anti-PSL2 LNAs to treat IAV well after an infection has been established, we sought to deliver our LNAs in a way most likely to be administered therapeutically to hospitalized, severely ill patients: by I.V. injection. Mice infected with a lethal dose of PR8 virus (n=10) were treated with either LNA14, LNA9, Scrambled LNA, or vehicle control by intravenous injection on day 3 post-infection (Day +3) when mice typically become noticeably ill, to simulate a severe, hospitalized infection. While mice treated with vehicle and scrambled controls rapidly succumbed to the infection, approximately 65% of mice treated with LNA9 survived lethal infection, and all mice in the LNA14-treated cohort survived (**Fig. 6m**).

## Discussion

We describe here the discovery and characterization of an RNA stemloop structure, PSL2, that serves as a packaging signal for genome segment PB2, which is conserved across influenza A isolates. Knowledge of PSL2’s RNA secondary structure not only can explain previous fortuitously discovered packaging-defective mutations, but also enabled the rational design of mutations with more potent disruption of packaging. Moreover, compensatory mutations that restore PSL2’s structure (but not primary sequence) rescue virus packaging and titer loss *in vitro*, thus providing strong genetic validation of the importance of PSL2’s RNA secondary structure for influenza virus packaging. Extending these findings *in vivo*, PSL2 disrupting and compensating mutants give striking attenuation and restoration, respectively, of mortality in mice—confirming PSL2’s importance not only to the viral life cycle, but also for influenza-mediated disease. Antisense locked nucleic acids (LNAs) designed to disrupt PSL2 structure dramatically inhibit IAV *in vitro* against viruses of different strains and subtypes, exhibit a high barrier to the development of resistance, and are equally effective against wild-type and NAI-resistant viruses. *In vivo,* intranasal dosing of LNAs resulted in potent antiviral efficacy and prevented mortality in mice, even with a single dose administered two weeks prior to infection with a lethal IAV inoculum. In a therapeutic model, an anti-PSL2 LNA provided complete protection from death when administered three days post-infection with a lethal IAV dose. Moreover, the PSL2-targeting LNAs not only provided full protection against lethal IAV infection, but also enabled the surviving mice to develop vigorous immunity. Together, these results have exciting implications for the development of a novel class of pan-genotypic anti-IAV therapies for prophylaxis of, treatment of established, and “just-in-time” universal vaccination against, an IAV infection.

Historically, it was assumed that IAV RNA secondary structures would be inaccessible in vRNP complexes due to the association between the vRNA and NP; however, local RNA structures can remain recognizable as substrates. For example, bound RNA in vRNP complexes are still accessible to treatment by RNases, and unprotected vRNA regions are sufficient for hybridization to complementary cDNA fragments^31,32^. Furthermore, some findings suggest the presence of NP may actually increase vRNA base accessibility and thus might enable intermolecular RNA-RNA interactions, rather than impede them^33^. We therefore hypothesized that local structural motifs may be exposed at the surface of the vRNPs, and could be enabling RNA-RNA or RNA-protein interactions that guide the packaging process.

This hypothesis led to the discovery of a novel stemloop structure, PSL2, in the 5′ terminal-coding region of IAV segment PB2 genomic vRNA known to be important for virus packaging. Previously-defined, as well as more informed and potent newly designed, packaging-defective mutations^13,16^ in this region destroyed PSL2 structure and yielded packaging-defective viruses that were attenuated *in vivo,* while compensatory mutations designed to restore the structure rescued virus packaging *in vitro* and restored lethality in an *in vivo* mouse model. Given the large body of work characterizing IAV packaging signals, we were initially surprised that the critical synonymous mutation sites, m52 and m65, had gone undiscovered or unremarked by prior studies. However, despite previous efforts that sequentially mutated the PSL2 region that included these nucleotide regions^13^, their mutational strategy suffered from lack of knowledge of the stemloop structure in two important ways. First, the previously described nt65 mutation was C65U, which still produces a basepaired G—U in the PSL2 stem, generating a structurally tolerated mutation. In contrast, our m65 (C65A) mutation does not base pair with the opposing guanine nucleotide and shows severe structure disruption (**Fig. 3, Supplementary Fig. 7**). Second, to our knowledge no previous work mutated the m52 nucleotide, but rather mutated its adjacent neighbor (nt C50A), which lies in an exposed bulge of PSL2. There are six possible permissible synonymous changes to the arginine amino acid (ARG 755) that covers nucleotide 52, almost all leading to a structurally permissible change. Without information about the PSL2 structure, this critical base pair and its relevance to PB2 packaging could easily be overlooked.

Importantly, we found that restoration of PSL2 structure through compensatory mutagenesis not only rescued packaging of its harboring segment PB2, but it also restored the incorporation of other segments. While genome segmentation provides a challenge to virus packaging, it confers an evolutionary advantage: namely, the ability for segments of different viral lineages and subtypes to be swapped during co-infection leading to the production of novel viruses. This propensity towards viral reassortment in both the form of antigenic drift, which leads to new strains that result in our need for yearly vaccine redesign, and viral shift, which introduces antigenically unique viral subtypes into the population that risk giving rise to new pandemics, necessitates a certain level of conservancy in the packaging signal regions of each segment to allow for reassortment between different viruses. The sequence region in segment PB2 that contains the PSL2 stemloop is highly conserved across IAV subtypes, strains, and host species restricted viruses, and likely reflects a strict biologic requirement for its preservation. SHAPE analysis of this region confirmed maintenance of the PSL2 structure between seasonal as well as pandemic viruses of different subtypes and host origins, suggesting that this structural element could be a novel broad-spectrum therapeutic target. Although our structural studies are based on *in vitro* RNA templates, we expect that rapidly emerging improvements in RNA structure probing technologies will enable a more detailed view of PSL2’s RNA structure within cells and in the context of the vRNP complex.

Recent studies have confirmed and further expanded on the role that RNA structure plays in the IAV lifecycle^34–43^. These include the findings that regions of low NP-binding fall within packaging signals and are likely to contain RNA structures^39,40^, the continued emergence of new RNA-RNA interactions between segments during the packaging process^34^, the discovery that specific nucleotide residues are important to the formation of precise vRNP structures critical to genome packaging^43^, the uncovering of new RNA secondary structures in the MX gene and their roles in the production of infectious particles^35,36^, and excitingly, the first global RNA structure of the IAV genome mapped in cells^41^, to name a few. As the technology to detect and monitor RNA structures and their interactions in the context of viral infection grows, we anticipate many new discoveries of the unique roles these structures, including PSL2, play in IAV and how they interact to guide the packaging and virus production process.

In our present study, we have demonstrated that LNA-mediated targeting of the PSL2 structure has potent antiviral activity *in vitro* across divergent IAV subtypes, and importantly, protects against lethal IAV infection *in vivo* after as little as a single intranasal dose given two weeks before infection, or three days post-infection. Moreover, mice treated with LNA prior to inoculation with a lethal IAV dose show robust long-lasting immunity against high titer virus rechallenge. One of the limits to the currently approved IAV vaccine strategy is its inadequate production of long-term humoral immunity, with recent research suggesting that viral infection is far superior to vaccination in the protective efficacy against heterosubtypic circulating virus^44–46^. Given the potential for long-lasting prophylactic protection after a single dose of LNA14, combined with the universality of the pan-genotypic PSL2 target, it is attractive to speculate on the clinical use of anti-PSL2 LNAs as a “just-in-time” universal vaccine strategy. In addition, because traditional vaccines take weeks to provide full protection, a co-administered single dose of our LNAs could provide protection during this vulnerability window.

The use of RNA-based therapeutics to treat disease is a rapidly growing field. The advent of LNA technology allows for the design of antisense oligonucleotides (ASO) that possess greater biologic capabilities than their siRNA and non-LNA base counterparts, both in terms of *in vivo* stability and target specificity. Several ASO-mediated therapeutics have now gained FDA-approval for use in humans^47,48^, and there are an ever-growing number of promising candidates currently in clinical trials^49,50^. Recently, an siRNA against respiratory syncytial virus in lung transplant recipients has shown success in a Phase 2b trial, where the siRNA was delivered nasally via a nebulizer^51^. We envision a similar method of treatment and path to the clinic for PSL2-targeted therapies, like LNA14 against IAV, although intravenous administration for severe hospitalized patients may be a complementary path. Moreover, analogous strategies can now be contemplated for rapidly identifying and targeting critical RNA secondary structures in a wide range of viruses for which no effective therapies currently exist. Importantly, incorporating the targeted virus’ RNA secondary structure into the design of antiviral LNAs allowed us to achieve far greater inhibition than with ASOs designed against the same viral genomic sequence but which relied only on primary nucleotide sequence homology for their design^52^.

Oseltamivir (Tamiflu™) is the most widely used and stockpiled neuraminidase inhibitor (NAI) on the market. Like all NAIs, OSLT requires a conformational rearrangement in the viral NA protein to accommodate the drug. Any mutations in the NA protein that affect this rearrangement reduce the binding affinity of OSLT, thus reducing drug efficacy. Notably, the H275Y mutation is most associated with OSLT resistance. Selection for resistance mutations is of particular concern with IAV, especially in immunocompromised populations, where it was shown that the rapid selection of the H275Y mutation in an immunocompromised patient lead to clinical failure of the last-resort NAI drug, peramivir, suggesting that the selection for multi-drug resistant viruses in immunocompromised hosts may be more common than previously believed^1^. Moreover, IAV has demonstrated the ability to acquire resistance to OSLT in unexpected ways^53^. This, together with the recent spread of OSLT-resistant and NAI-resistant viruses in circulation^54,55^, indicates that we may need to reevaluate our usage of NAIs as a whole and are in urgent need of orthogonal anti-IAV therapeutics.

Baloxavir marboxil (“BXM”, Xofluza™), a new-in-class antiviral agent that inhibits IAV replication by targeting the endonuclease function of the viral polymerase complex, was approved by the FDA in late 2018. It is the first novel flu treatment to receive FDA approval since the clearance of NAIs oseltamivir and zanamivir in 1999^56^. Like NAIs, however, BXM targets a viral protein, making the development of drug resistance a concern. Indeed, in its first year of use, reports of viruses with reduced susceptibilities to BXM were isolated from cell cultures, from patients in clinical trials, and from adults infected with A/H1N1, as well as pediatric patients infected with A/H3N2^57,58^. The rapid emergence of resistance mutations and the ease at which they arise in both the circulating virus subtypes, constitute a clear warning for the drug’s clinical efficacy. Here, we demonstrate that PSL2-targeted LNAs can possess a high barrier to the development of antiviral resistance, and that LNAs can retain their same high potency against NAI-resistant strains. Such discovery of new viral targets and the development of new classes of antivirals is imperative to reduce the adverse impact current and future pandemics can have on human health.

## Acknowledgements

We thank Hong Jin and her team at MedImmune for the kind donation of chicken eggs and technical guidance. Additional thanks go to Phil Pang for initial study brainstorming. The work was supported in part by a Mona M. Burgess Stanford BIO-X Interdisciplinary Graduate Fellowship, the National Institutes of Health (NIH) Graduate Training Grant 5T32AI007328-24, NIH research grants R56A1111460, U19A1109662, RO1AI132191, a Harrington Scholar Innovator Grant, and W81XWH1810647 from USAMRAA, Department of Defense. The work was also supported in part by the Intramural Research Program of the National Institute of Allergy and Infectious Diseases, National Institutes of Health. The data presented in this manuscript are tabulated in the main paper and in the supplementary materials. Chemical mapping datasets for mutate-and-map and mutation/rescue experiments have been deposited at the RNA Mapping Database (http://rmdb.stanford.edu). R.J.H. and J.S.G. conceived and designed the study. R.J.H., M.E., L.B., K.N., A.X. and S.T. performed the experiments. R.J.H., M.E., S.T., and L.B. analyzed the data. M.E. and T.A. provided expertise and assistance for mouse experiments. B.F. performed sequencing and assisted in experimental prep. P.L. assisted in cloning of constructs. E.A.P. aided in designing oligonucleotides, experimental planning, results analysis, and compilation of manuscript. W.K. provided important technical assistance on SHAPE experiments. S.S. and P.K. provided statistical analyses. J.K.T. offered valuable expertise and coordinates before publication, discussed results, and provided critical reagents. R.D. and J.S.G. provided experimental feedback and assisted in results analysis. R.J.H., J.S.G., M.E., S.T. and R.D. wrote the paper. The authors declare competing financial interests: A patent pertaining to the materials presented in this article has been filed with the U.S. Patent and Trademark Office.

## Materials and Methods

### Cells and viruses

HEK 293T and MDCK cells (NLB-2) were obtained from American Type Culture Collection ‘ATCC’ (Manasass, VA) and were maintained in Dulbecco’s modified Eagle’s medium with 10% fetal bovine serum and penicillin-streptomycin (Gibco). All cell lines used in this report were routinely checked for mycoplasma contamination (MycoAlert Mycoplasma Detection Kit, Lonza) and were authenticated by the respective vendors. Wild-type influenza A/PR/8/34 (PR8) H1N1 virus (ATCC-VR-95) and the tissue-culture adapted PR8 virus (ATCC-VR-1469) were purchased from ATCC. PR8 mutant viruses were generated using an eight-plasmid reverse genetic system as previously described^59^. Tissue-cultured adapted influenza A/Hong Kong/8/68 (HK68) H3N2 virus (ATCC-VR-1679), A/Virginia/ATCC6/2012 (H3N2) virus (ATCC-VR-1811), A/Virginia/ATCC1/2009TC (H1N1) virus (ATCC-VR-1736), and A/Wisconsin/33 (H1N1) virus (VR-1520) were purchased from ATCC. A/California/4/2009 (pH1N1) virus was kindly gifted by Elena Govorkova from St. Jude Children’s Research Hospital (Memphis, USA). Viruses were grown and amplified in 10-day-old specific-pathogen-free research grade chicken embryos at 35°C (Charles River Laboratories; SPAEAS).

### Plasmid constructs and cloning

Plasmids were used containing the wild-type PB2 segments from influenza viruses A/PuertoRico/8/34 (H1N1) [PR8], A/New York/470/2004 (H3N2) [NY470], A/New York/312/2001 (H1N1) [NY312], A/Brevig Mission/1/1918 (H1N1) [1918], A/California/04/2009 (H1N1) [CA09], and A/Vietnam/03/2004 (H5N1) [VN1203]. For the generation of PR8 packaging mutant vRNA, we utilized a Stratagene QuickChange XL site-directed mutagenesis kit (Stratagene) for mutagenesis of a pDZ plasmid containing the PB2 gene of PR8^59^. Sequences of each mutated construct were confirmed by automated sequencing. The 8-plasmid pBD rescue system for A/WSN/33 (H1N1) was kindly donated by Andrew Mehle. The H275Y NA mutant was generated by QuickChange mutagenesis from the bidirectional pBD plasmids, as described above.

### Reverse genetics and virus titrations

Recombinant A/Puerto Rico/8/34 (PR8) virus and recombinant A/WSN/33 (WSN) virus were generated using eight-plasmid reverse genetic systems^59^. Briefly, 10^6^ cells of a 293T/MDCK co-culture were Lipofectamine™ 3000 (Invitrogen™) transfected with 1μg of one of each of the eight segments contained within plasmids that utilize a bidirectional dual Pol I/II promoter system for the simultaneous synthesis of genomic vRNA and mRNA. For rescue of compensatory PB2 mutant viruses where a non-synonymous change was required, a wild-type PB2 protein expression plasmid (Pol II) was co-transfected during virus rescue. Supernatants were collected 24 hours post-transfection. PR8 rescue viruses were then inoculated into the allantoic cavities of 10-day-old chicken embryos. WSN rescue viruses were passaged subsequent times on MDCK cells. Rescue of recombinant viruses was assessed by hemagglutination activity. Each newly rescued virus was further plaque titered and mutations were confirmed by sequencing of mutated genes. Plaque assays were carried out on confluent MDCK cells as described previously^60^. Hemagglutination (HA) assays were carried out in 96-well round-bottomed plates at room-temperature, using 50 μl of virus dilution and 50 μl of a 0.5% suspensions of turkey red blood cells (LAMPIRE^®^ Biological Laboratories) in phosphate-buffered saline (PBS).

### Isolation of packaged vRNAs

To analyze packaged vRNA for PR8 mutated viruses, 10-day-old eggs were inoculated with approximately 1000 PFU of recombinant virus and incubated for 72 hours. Allantoic fluid was harvested, and supernatant was dual-clarified by low-speed centrifugation. Clarified supernatant was then layered on a 30% sucrose cushion and ultra-centrifuged at 30,000 RPM for 2.5 hours (Beckman Rotor SW41). Pelleted virus was resuspended in PBS and TRIzol (Invitrogen) extracted. Precipitated vRNA was resuspended in a final volume of 20 μl of 10mM Tris-HCl (pH 8.0) and stored at −80°C.

Virus supernatant from LNA-treated cells was harvested 48hpi and subjected to low-speed centrifugation at 1000 RPM then 10,000 RPM. Isolation continued as indicated above.

### qPCR analysis of packaged vRNAs

Approximately 200 ng of extracted vRNA was reverse transcribed using a universal 3′ primer (5′-AGGGCTCTTCGGCCAGCRAAAGCAGG) and Superscript III reverse transcriptase (RT) (Invitrogen). The RT product was diluted approximately 10,000-fold and used as a template for quantitative PCR (qPCR). Separate PCRs were then carried out as previously described^61^ with segment-specific primers. The 10 μl reaction mixture contained 1 μl of diluted RT product, a 0.5 μM primer concentration, and SYBR Select Master Mix (Applied Biosystems) that included SYBR GreenER dye, 200 μM deoxynucleoside triphosphates, heat labile UDG, optimized SYBR Green Select Buffer, and AmpliTaq DNA polymerase UP enzyme. Relative vRNA concentrations were determined by analysis of cycle threshold values, total vRNA amount within a sample was normalized to the level of HA vRNA, and then percentages of incorporation were calculated relative to the levels of wild-type vRNA packaging. Viral packaging results represent the averaged levels of vRNA incorporation ± standard deviations derived from two independent virus purifications, with vRNA levels quantified in triplicate.

### Strand-specific RT-qPCR

MDCK cells transfected with 1 mM LNA-9 or scrambled LNA were infected with PR8 virus at an MOI of 0.1 24 hours post transfection. Eight hours post infection total cellular RNA was extracted in Trizol reagent (Invitrogen) and the RNA was purified using the Direct-Zol RNA mini-prep (Zymo Research) according to the manufacturer protocol. Reverse transcription and qPCR were performed according to^62^. cDNAs of the influenza viral RNA (vRNA) and complementary viral RNA (cRNA) were synthesized with tagged primers to add an 18–20 nucleotide tag that was unrelated to the influenza virus at the 5′ end (cRNAtag; 5′-GCT AGC TTC AGC TAG GCA TC-3’, vRNAtag; 5’-GGC CGT CAT GGT GGC GAA T-3′). Hot-start reverse transcription with the tagged primer was performed as described in Kawakami et al., 2011 using saturated trehalose. A 5.5 μl mixture containing 200 ng of total RNA sample and 10 pmol of tagged primer were heated for 10 min at 65°C, chilled immediately on ice for 5 min, and then heated again at 60°C. After 5 min, 14.5 μl of preheated reaction mixture [4 μl First Strand buffer (5 ×, Invitrogen), 1 μl 0.1 M dithiothreitol, 1 μl dNTP mix (10 mM each), 1 μl Superscript III reverse transcriptase (200 U/μl, Invitrogen), 1 μl RNasin Plus RNase inhibitor (40 U/μl, Promega) and 6.5 μl saturated trehalose] was added and incubated for 1 h. Real-time PCR (qPCR) was performed with PowerUp SYBR Green SuperMix (Applied Biosystems) on a BIORAD CFX96 Real-Time System. Seven microliters of a ten-fold dilution of the cDNA was added to the qPCR reaction mixture [10 μl SYBR Green SuperMix (2 ×), 1.5 μl forward primer (10 μM), 1.5 μl reverse primer (10 μM)]. The cycle conditions of qPCR were 95°C 10 min, followed by 40 cycles of 95°C 15 sec and 60°C for 1 min. qPCR primers were: PR8 segment 1 (PB2) cRNA, Forward: 5′-TCC ACC AAA GCA AAG TAG AAT GC-3′; Reverse: 5′-GCT AGC TTC AGC TAG GCA TCA GTA GAA ACA AGG TCG TTT TTA AAC-3′. PR8 segment 1(PB2) vRNA, Forward: 5′-GGC CGT CAT GGT GGC GAA TAG ACG AAC AGT CGA TTG CCG AAG C-3′, Reverse: 5′-AGT ACT CAT CTA CAC CCA TTT TGC-3′. PR8 segment 4 (HA) cRNA, Forward: 5′-CTG TAT GAG AAA GTA AAA AGC C-3’, Reverse: 5′-GCT AGC TTC AGC TAG GCA TCA GTA GAA ACA AGG GTG TTT TTC-3′. PR8 segment 4 (HA) vRNA, Forward: 5′-GGC CGT CAT GGT GGC GAA TAG GAT GAA CTA TTA CTG GAC CTT GC-3′, Reverse: 5′-TCC TGT AAC CAT CCT CAA TTT GGC-3′.

### Animals

All animal studies were performed in accordance with the National Institutes of Health Guidelines for the Care and Use of Laboratory Animals and approved by the Stanford University Administrative Panel on Laboratory Animal Care. Six to eight Healthy age-matched female BALB/c mice (Jackson Laboratories, Bar Harbor ME) were randomly separated into groups for infection/treatment or used as uninfected/non-treated controls. Treatment groups were not blinded to the investigators. Mice were identified with tag numbers throughout the experiment.

### *In vivo* infection

Mice were lightly anesthetized with isoflurane and intranasally infected with with 50 μl of virus preparation at a concentration of 1000 PFU for virus packaging mutant experiments and 900 PFU for LNA treatment experiments. Weights and clinical scores were assessed daily, and animals were humanely sacrificed when a clinical score of 5 was recorded (*see* Supplementary Table 2 for clinical score determination). Kaplan-Meier survival curves were generated using GraphPad Prism.

### *In vivo* antiviral assays

*‘In vivo*-ready’ LNAs were custom designed and ordered from Qiagen (formally Exiqon). For intranasal delivery, *in vivo-ready* LNA was mixed in complexes with *In vivo-*JetPEI® transfection reagent (Polyplus) according to manufacturer’s protocol to the indicated final concentration in 50 μl of 5 % glucose solution. Mice were then lightly anesthetized with isoflurane and 50 μl of the solution was delivered intranasally. For retro-orbital delivery: *‘In vivo*-ready’ LNA was mixed in complexes with *In vivo-*JetPEI® transfection reagent (Polyplus) according to manufacturer’s protocol to the indicated final concentration in 200 μl of 5 % glucose solution. Mice were then anesthetized, and the solution was delivered by retro-orbital injection.

### Locked Nucleic Acid (LNA) design and preparation

Oligonucleotides containing locked nucleic acids (LNA) were custom synthesized from Exiqon (Vedbaek, Denmark), and later by IDT. Capitalized letters denote LNA. Lowercase letters denote typical (non-locked) DNA nucleotides. All oligonucleotides contain phosphorothioate internucleoside linkages. LNA 8 and 9 were designed as LNA gapmers to contain a stretch of 6-7 DNA nucleotides optimized for RNAse-H recruitment. Sequences of all LNAs are shown below.

**LNA 1: 5′** AccAaaAGaaT **3′**
**LNA 2: 5′** TggCcATcaaT **3′**
**LNA 3: 5′** TagCAtActtA **3′**
**LNA 4: 5′** CCAAAAGA **3′**
**LNA 5: 5′** CATACTTA **3′**
**LNA 6: 5′** CagaCaCGaCCaaAA **3′**
**LNA 7: 5′** TAcTtaCTgaCagCC **3′**
**LNA 8: 5′** AGAcacgaccaaAAG **3′** –with RNase-H activity
**LNA 9: 5′** TACTtactgacaGCC **3′** –with RNase-H activity
**LNA14: 5′** CGACcaaaagaATTC **3′** –with RNase-H activity
**Scramble LNA (negative control): 5’** AACACGTCTATACGC **3’**

### *In vitro* LNA antiviral assays

LNAs were reconstituted in RNAse-free water at 100 μM stock solutions, aliquoted and stored at −20 °C prior to single-use. Lipofectamine 3000® (Life Technologies) was used to transfect LNA into cells at indicated concentrations per manufacturer’s protocol. For prophylactic antiviral assays, 10^6^ MDCK cells were plated in 6-well plates 24 hours prior to being transfected with the indicated LNA. Cells were then infected at the indicated time points with 0.01 MOI of PR8 (H1N1) or HK68 (H3N2) virus. For post-infection therapeutic antiviral assessment, MDCK cells were infected with PR8 or HK68 prior to LNA transfection as described above. Forty-eight hours post-infection, supernatant was collected, and viral titer was determined by plaque assay in triplicate.

### LNA-treatment and packaging efficiency determination

Briefly, T75 flasks of 80% confluent MDCK cells were transfected with 100 nM of scrambled LNA, LNA9, or mock untreated by Lipofectamine 3000 transfection, according to manufacturer’s protocol. Twelve hours post transfection, cells were infected with 0.01 MOI of wild-type TC-adapted PR8 virus. After 1 hour, virus was removed and cells were washed with PBS. Forty-eight hours post-infection supernatants were collected, and RNA was isolated as described in isolation of packaged vRNAs and assay methods.

### *In vitro* drug selection

LNA9 selection: 80-90% confluent MDCK cells in 12-well plates were transfected in duplicate with a starting concentration of 0.01 nM (∼1/2 EC_50_) LNA9 for Passage 1 by Lipofectamine transfection (see above). Twelve hours post-transfection, cells were washed with PBS and infected with 0.01 MOI of wild-type PR8 virus. After 1 hour incubation at 37 °C, cells were washed and virus growth media was added. Cells were incubated until 50 % CPE was evident (48-72 hours). Virus supernatant was harvested, low-speed centrifuge clarified, aliquoted, plaque titered and stored at −80°C. The virus supernatant was then continuously serially passaged in the presence of escalating concentrations of LNA9 (0.01 nM to 100 nM). If no CPE was evident, drug concentration was lowered and added virus concentration was increased until 50 % CPE occurred. OSLT selection: confluent MDCK cells in 12-well plates were infected with 0.01 MOI of PR8 virus. After adsorption for an hour, cells were washed with PBS, and OSLT (Sigma Aldrich Cat. No. Y0001340) was added to virus growth media at a starting concentration of 1 nM (∼1/2 EC_50_). Drug selection proceeded as described above, with escalating concentrations of OSLT (0.01 nM to 250 μM) at each subsequent passage.

### EC_50_ determination

For LNA9, the 50% effective concentration (EC_50_) was defined as the concentration of drug effective in reducing the percent of virus titer to 50% of that for the no-drug control. In brief, the EC_50_ was determined by seeding 5 × 10^5^ MDCK cells in each well of a 12-well plate and incubating overnight at 37°C under 5% CO_2_. Cells were then transfected with LNA9 as described above at concentrations from 0.01 nM to 10 μM. Plates were incubated at 37°C for 12 hours prior to infection with 0.01 MOI of wild-type PR8, serially passaged LNA-treated virus, WSN33 wild-type or WSN33 H275Y NAI-resistant virus. Forty-eight hours post-infection, supernatants were collected, centrifuge clarified, aliquoted and stored at −80°C. The viral titer for each drug dilution was performed by plaque assay in duplicate. The EC_50_ was the concentration of LNA9 yielding a percent titer of 50% of that without drug.

For OSLT, the EC_50_ was defined as the concentration of drug reducing the total percentage of plaques to 50% of that for the no-drug control, determined by plaque reduction assay^1^. Briefly, confluent MDCK cells in 12-well plates were infected with approximately 100 PFU of wild-type PR8, serially passaged OSLT-treated virus, WSN33 wild-type or WSN33 H275Y NAI-resistant virus and incubated for 1 hour at 37 °C. Cells were then washed with PBS, and a 50:50 mix of 1 % agarose to 2x virus growth DMEM containing varying concentrations of drug (0.1 nM to 1 mM) was added to the cells. Plates were harvested 72 hours later, stained with crystal violet, and plaques were counted. The EC_50_ was the concentration of OSLT reducing the total percentage of plaques to 50 % of that without drug. All results were plotted in GraphPad Prism to generate EC_50_ curves.

### *In vitro* transcription of full-length vRNA

For each wild-type isolate (PR8, 1918, VN1203, NY470, NY312, and CA09) and PR8 packaging mutant clones, PB2 cDNA was amplified from plasmid using segment-specific primers under a T7 promoter. Amplified cDNA was gel-purified using an Invitrogen DNA gel kit. vRNAs were then produced by *in vitro* transcription, using T7-MEGAscript kit. vRNAs for SHAPE were purified by MEGAclear (Thermofisher, cat. no. AM1908) with purity and length verified by capillary electrophoresis.

### sf-SHAPE analysis of full-length IAV vRNA

*In vitro* transcribed PB2 vRNA was folded (100 mM NaCl; 2.5 mM MgCl; 65 °C x 1’, 5′ cooling at room temperature, 37 °C for 20–30’) in 100 mM HEPES, pH=8. 2’ acylation with NMIA^19^ and reverse transcription (RT) primer extension were performed at 45 °Cx 1’, 52 °C x 25’, 65 °C x 5’, as previously described^63^. 6FAM was used for all labeled primers (primer sequences available upon request). Exceptions to these protocols were as follows: (i) RNA purification after acylation was performed using RNA C&C columns (Zymo Research), rather than ethanol precipitation; (ii) before and after SHAPE primer buffer was added, the mixture was placed at room temperature for 2–5 min, which enhanced RT transcription yields significantly; (iii) DNA purification was performed using Sephadex G-50 size exclusion resin in 96-well format then concentrated by vacuum centrifugation, resulting in a more significant removal of primer; and (iv) 2 pmol RNA was used in ddGTP RNA sequencing reactions.

The ABI 3100 Genetic Analyzer (50 cm capillaries filled with POP6 matrix) was set to the following parameters: voltage 15 kV, T = 60°C, injection time=15 s. The GeneScan program was used to acquire the data for each sample, which consisted of purified DNA resuspended in 9.75 μl of Hi-Di formamide, to which 0.25 μl of ROX 500 internal size standard (ABI Cat. 602912) was added. *PeakScanner* parameters were set to the following parameters: smoothing=none; window size=25; size calling=local southern; baseline window=51; peak threshold=15. Fragments 250 and 340 were computationally excluded from the ROX500 standard^64^. The data from *PeakScanner* were then processed into SHAPE data by using FAST (fast analysis of SHAPE traces), a custom algorithm developed in our lab^20^. FAST automatically corrects for signal differences due to handling errors, adjusts for signal decay, and converts fragment length to nucleotide position, using a ddGTP ladder as an external sizing standard and the local Southern method ^20,65^. This algorithm embedded in the *RNAstructure program* is freely available at http://med.stanford.edu/glennlab/download.html.

*RNAstructure* parameters: slope and intercept parameters of 2.6 and −0.8 kcal/mol, were initially tried, as suggested^66^; however, we found that smaller intercepts closer to 0.0 kcal/mol (e.g. ∼ −0.3) produced fewer less optimal structures (within a maximum energy difference of 10%). We speculate that this minor parameter difference may be due to the precise fitting achieved between experimental and control data sets by the automated FAST algorithm. FAST was written in ANSI C/C++ and is integrated into *RNAstructure* with FAST, which requires MFC (Microsoft Foundation Classes). RNA structures were drawn and colored using RNAViz 2^67^ and finalized in Adobe Illustrator.

### PSL2 Construct design, RNA synthesis and chemical modification for Mutate-and-Map Experiments

Double-stranded DNA templates were prepared by PCR assembly of DNA oligomers designed by an automated MATLAB script as previously described (NA_Thermo, available at https://github.com/DasLab/NA_thermo)68. Constructs for mutate-and-map (M^2^) includes all single mutants to Watson-Crick counterpart. Compensatory mutants for mutation/rescue were designed based on base-pairing in the proposed secondary structure^23^. *In vitro* transcription reactions, RNA purification and quantification steps were as described previously^68^. One-dimensional chemical mapping, mutate-and-map (M^2^), and mutation/rescue were carried out in 96-well format as described previously^68–70^. Briefly, RNA was heated up and cooled to remove secondary structure heterogeneity; then folded properly and incubated with SHAPE reagent (5 mg/mL 1-methyl-7-nitroisatoic anhydride (1M7))^71^; modification reaction was quenched and RNA were recovered by poly(dT) magnetic beads (Ambion) and FAM-labeled Tail2-A20 primer; RNA was washed by 70% ethanol (EtOH) twice and resuspended in ddH_2_O; followed by reverse transcription to cDNA and heated NaOH treatment to remove RNA. Final cDNA library was recovered by magnetic bead separation, rinsed, eluted in Hi-Di formamide (Applied Biosystems) with ROX-350 ladder, loaded to capillary electrophoresis sequencer (ABI3100). Data processing, structural modeling, and data deposition: The HiTRACE software package version 2.0 was used to analyze CE data (both MATLAB toolbox and web server available^72,73^). Trace alignment, baseline subtraction, sequence assignment, profile fitting, attenuation correction and normalization were accomplished as previously described^74,75^. Sequence assignment was accomplished manually with verification from sequencing ladders. Data-driven secondary structure models were obtained using the Fold program of the *RNAstructure* package version 5.4^76^ with pseudo-energy slope and intercept parameters of 2.6 kcal/mol and −0.8 kcal/mol. 2-dimensional Z score matrices for M^2^ datasets, and helix-wise bootstrapping confidence values were calculated as described previously^23,68^. Z score matrices were used as base-pair-wise pseudofree energies with a slope and intercept of 1.0 kcal/mol and 0 kcal/mol. Secondary structure images were generated by VARNA^77^. All chemical mapping datasets, including one-dimensional mapping, mutate-and-map, and mutation/rescue, have been deposited at the RNA Mapping Database (http://rmdb.stanford.edu)78, accession codes: PSL2IAV_1M7_0001, PSL2IAV_RSQ_0001.

### SHAPE analysis of LNA-targeted vRNA

A truncated DNA template of PR8 virus segment PB2 containing nucleotides 1-88 was prepared by PCR assembly of DNA oligomers, and *in vitro* transcription reactions, RNA purification and quantification steps were as described previously^68^. One-dimensional SHAPE chemical mapping was performed in 96-well plate format as described above with the following exception: once RNA was denatured and refolded as described, 100 nM of each prepared LNA was added to the folded RNA and incubated with 5 mg/mL of SHAPE reagent 1M7 (1-methyl-7-nitroisatoic anhydride). Modification quenching, RNA recovery, re-suspension, reverse transcription, cDNA sequencing and data processing were performed as described in ref. 44.

### Statistical analyses

We expressed the data as the mean ± s.d. We used Student’s *t*-test (to compare two samples) or ANOVA (to compare multiple samples) (GraphPad InStat 3) for statistical analysis. We performed the Kaplan-Meier log-rank test for survival analyses. We considered all *P* values >0.05 not to be significant.

