## Supplementary Materials for "Identification and targeting of a pan-genotypic influenza A virus RNA structure that mediates packaging and disease"

**This PDF file includes:**

**Supplementary Fig. 1.** SHAPE reactivity and nucleotide conservancy of PB2 vRNA
**Supplementary Fig. 2.** Packaging-defective mutations disrupt wild-type SHAPE reactivity
**Supplementary Fig. 3.** 2-Dimensional Mutate-and-Map (M2) analysis of PSL2 RNA secondary structure
**Supplementary Fig. 4.** Synonymous mutation of single highly conserved codons of the PR8 PB2 vRNA
**Supplementary Fig. 5.** Effect of synonymous mutation on PSL2 structure
**Supplementary Fig. 6.** Effect of compensatory mutations in PR8 PB2 packaging-defective mutants on viral packaging and titer
**Supplementary Fig. 7.** Mutate-Map-Rescue ($M^2R$) analysis
**Supplementary Fig. 8.** Multi-cycle growth curve of packaging-defective mutant viruses
**Supplementary Fig. 9.** Design of primer sequences for 2-dimensional M2R mutants
**Supplementary Fig. 10.** Replicon assay of defective and compensatory PB2 packaging mutants
**Supplementary Fig. 11.** Western blot of mutant PB2 protein expression in virus infected cells
**Supplementary Fig. 12.** SHAPE analysis on LNA-RNA interaction
**Supplementary Fig. 13.** Immunofluorescence analysis of LNA9 and vRNP in LNA treated infected cells
**Supplementary Fig. 14.** Western blot analysis of LNA9 and scrambled LNA treated infected cells
**Supplementary Fig. 15.** qPCR analysis of vRNA and cRNA in LNA9 and scrambled LNA treated infected cells
**Supplementary Fig. 16.** LNA9 treatment of cells has no cellular cytotoxicity
**Supplementary Fig. 17.** LNA9 treatment of cells infected at high MOI
**Supplementary Fig. 18**. Lung virus titers of LNA-treated mice post-infection

**Supplementary Table 1.** PB2 Packaging mutant nomenclature and corresponding sites of mutation
**Supplementary Table 2.** Description of clinical scoring in diseased mice

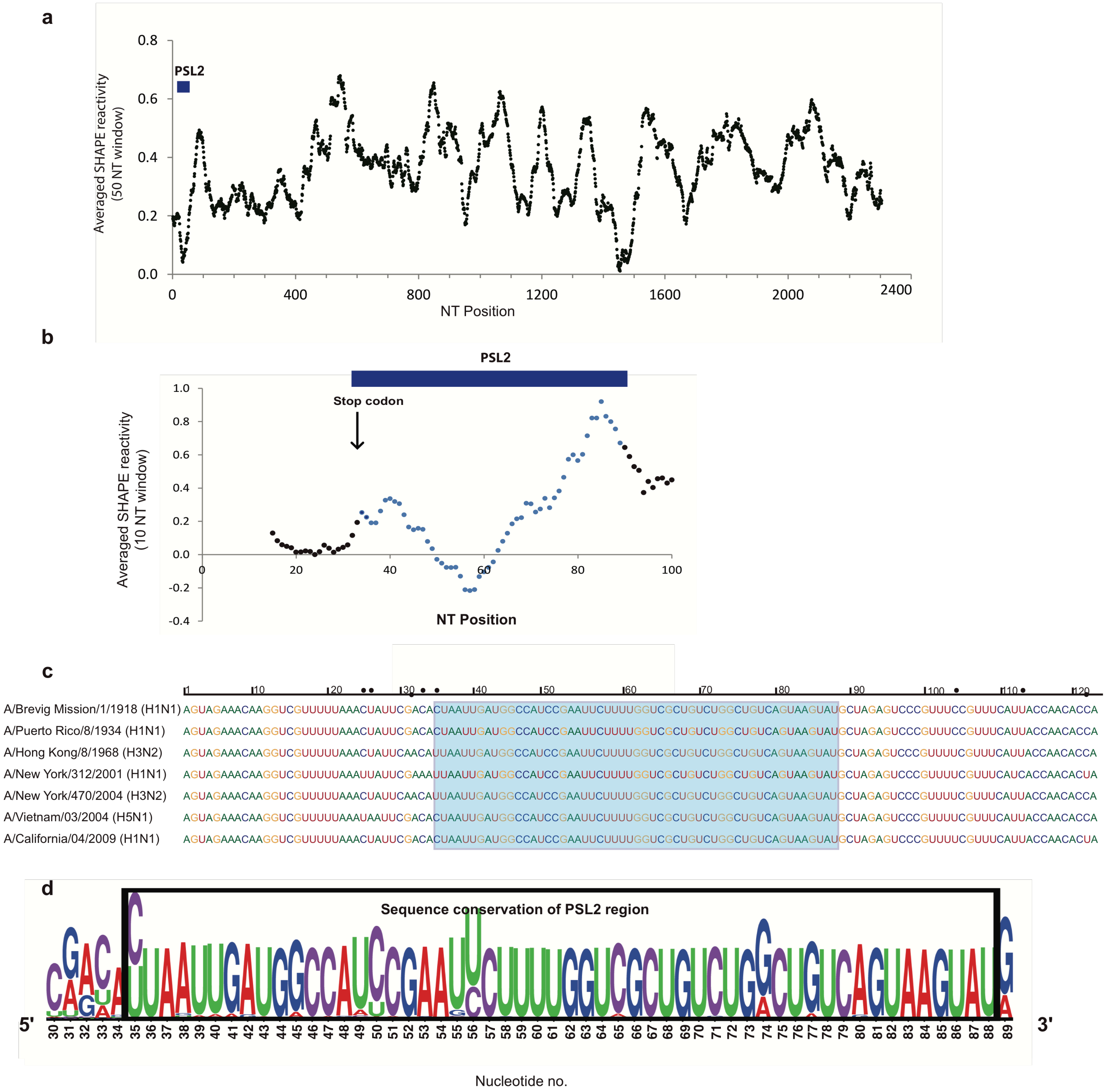


**Supplementary Figure 1**. **SHAPE reactivity and nucleotide conservancy of PB2 vRNA**. (**a**) Average SHAPE reactivity (window bin size = 100 nt) as a function of nucleotide position for the full-length (-)-sense PB2 vRNA from IAV strain A/Puerto Rico/8/1934 (H1N1). The PSL2 region (highlighted in blue, nts 34-86) encompasses the 5′ packaging signal domain, which possesses a high density of codons whose third position is conserved, and has one of the lowest SHAPE reactivities within the vRNA. The region after PSL2 is relatively unstructured yet contains another potential site for the presence of RNA structure between nucleotides 1400 and 1500. Interestingly, this second internal region was also predicted to contain structural elements by a 2011 bioinformatics study (ref. 17). (**b**) Zoomed in view of PSL2 region from (A). Window bin size = 10 nt. (**c**) Sequence alignment of the terminal 5′ region of PB2 corresponding to the following representative sequences from Figure 1d: pandemic A/Brevig Mission/1/1918 (H1N1), pandemic A/California/04/2009 (H1N1), seasonal human A/New York/470/2004 (H3N2), seasonal human A/Puerto Rico/8/1934 (H1N1), high pathogenic avian A/Vietnam/03/2004 (H5N1), pandemic A/Hong Kong/8/1968 (H3N2), and seasonal human A/New York/312/2001 (H1N1). RNA nucleotides are numbered in (-)-sense vRNA orientation. Shaded blue box encompasses the PSL2 RNA secondary structural element sequence. Black dots designate divergent nucleotide sites. (**d**) Graphical representation of nucleic acid sequence alignments across diverse influenza A viral subtypes and strains. Multiple sequence alignment was performed using MAFFT (Version 7) on 33,375 IAV PB2 sequences available in GenBank and visualized as a sequence logo (weblogo.berkeley.edu). The overall height represents sequence conservation at that nucleotide position, while the height of symbols within each position indicates the relative frequency of each nucleotide at that site. Black box = PSL2 region.

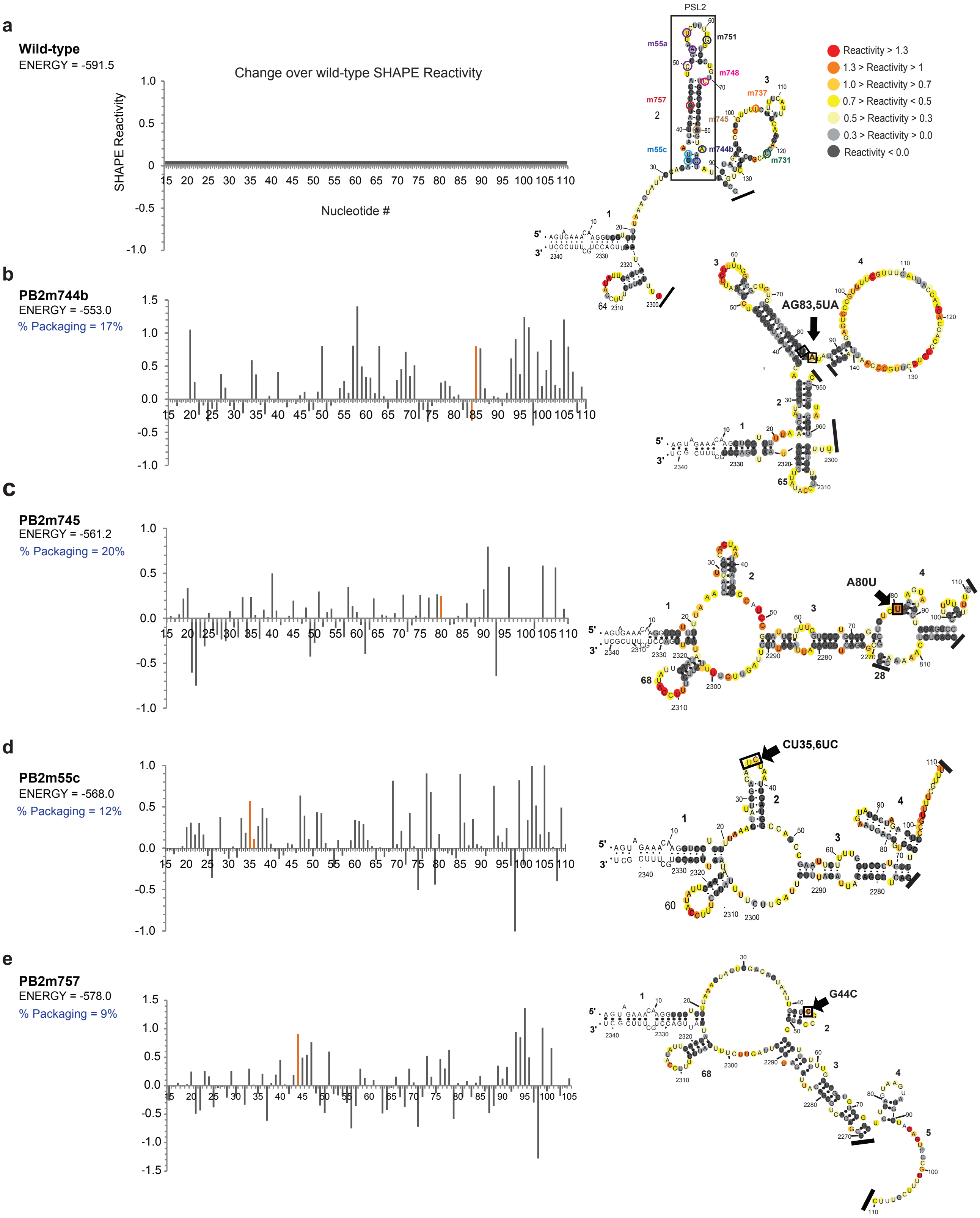
a
Wild-type
ENERGY = -591.5
Change over wild-type SHAPE Reactivity
SHAPE Reactivity
Nucleotide #
PSL2
m55a
m751
m748
m757
m745
m737
m55c
m744b
m731
Reactivity > 1.3
1.3 > Reactivity > 1
1.0 > Reactivity > 0.7
0.7 > Reactivity < 0.5
0.5 > Reactivity > 0.3
0.3 > Reactivity > 0.0
Reactivity < 0.0
b
PB2m744b
ENERGY = -553.0
% Packaging = 17%
AG83,5UA
c
PB2m745
ENERGY = -561.2
% Packaging = 20%
A80U
d
PB2m55c
ENERGY = -568.0
% Packaging = 12%
CU35,6UC
e
PB2m757
ENERGY = -578.0
% Packaging = 9%
G44C

**Supplementary Figure 2. Packaging-defective mutations disrupt wild-type SHAPE reactivity.** Left: Mutant SHAPE reactivity plotted as change over WT. Nucleotide numbering starts from 5′ end of (-)-sense vRNA. Orange bars indicate site of mutation(s). Energy values represent ΔG free energy of the predicted structure generated by the *RNAstructure* modeling algorithm using SHAPE pseudofree energy parameters. Right: SHAPE-determined structures of full-length (-)-sense mutant PB2 vRNA from PR8 strain (H1N1). Images are truncated to highlight 5′ terminal region. (**a**) Wild-type. (**b**-**e**) Packaging-defective mutants: (**b**) m744b (AG83, 85UA). (**c**) m745 (A80U). (**d**) m55c (CU35, 36UC). (**e**) m757 (G44C).

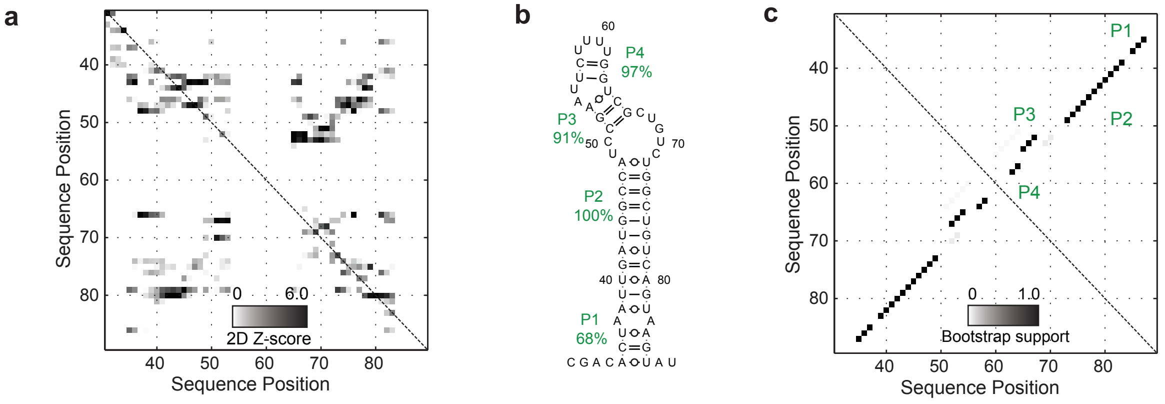


**Supplementary Figure 3. 2-Dimensional Mutate-and-Map (M2) analysis of PSL2 RNA secondary structure.** (**a**) Strong features of mutate-and-map data isolated by Z-score analysis. Z-scores were calculated for each nucleotide reactivity by subtracting the average reactivity of this nucleotide across all mutants and dividing by standard deviation (output_Zscore_from_rdat in HiTRACE). Squares show secondary structure model guided by mutate-and-map data. Dark signals highlight evidence of structured nucleotide pairing. (**b**) RNA secondary structure for 5′ packaging signal region (nts 30 – 88) derived from incorporating Z-scores into the *RNAstructure* modelling algorithm: bootstrap confidence estimates given as green percentage values. (**c**) Bootstrap support values for each base-pair shown as grayscale shading. Bootstrap values provide numerically accurate indicators of structural confidence. Low bootstrap confidence values suggest existence of alternative structural models.

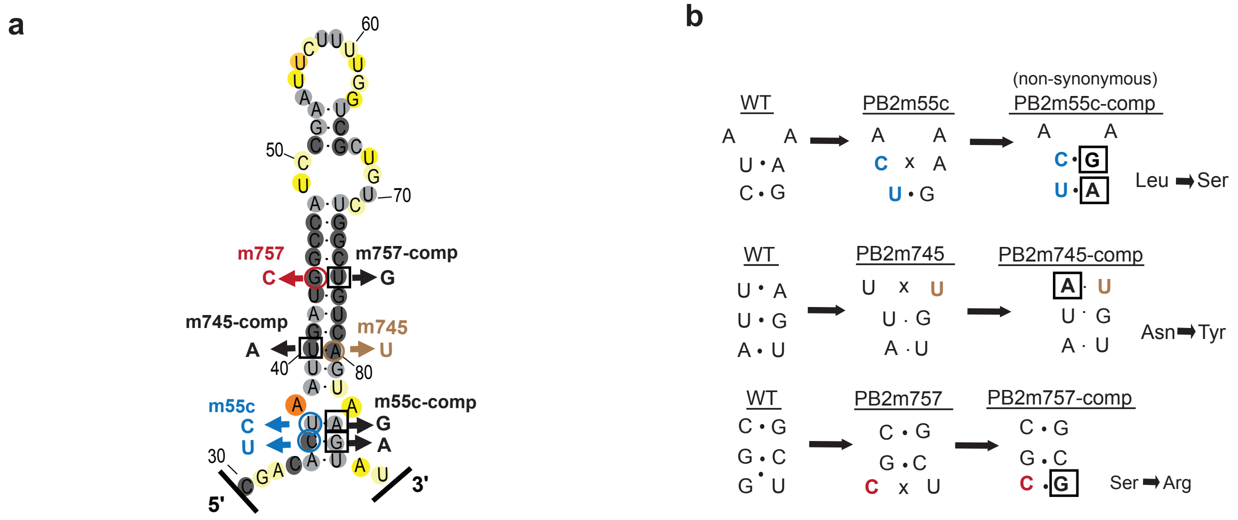


**Supplementary Figure 4. Synonymous mutation of single highly conserved codons of the PR8 PB2 vRNA.** (**a**) Previously described synonymous mutants (m757, m745, m55c) are mapped onto PSL2 structure. (**b**) Compensatory mutations (m55c-comp, m745-comp, and m757-comp) were designed at sites predicted to restore wild-type PSL2 structure based on SHAPE and mutate-and-map chemical analyses. Black boxed nucleotides denote site of compensatory mutation. (-)-sense vRNA orientation is shown. For mutations where a non-synonymous change was required to restore the structure, the alteration in encoded protein sequence is indicated.

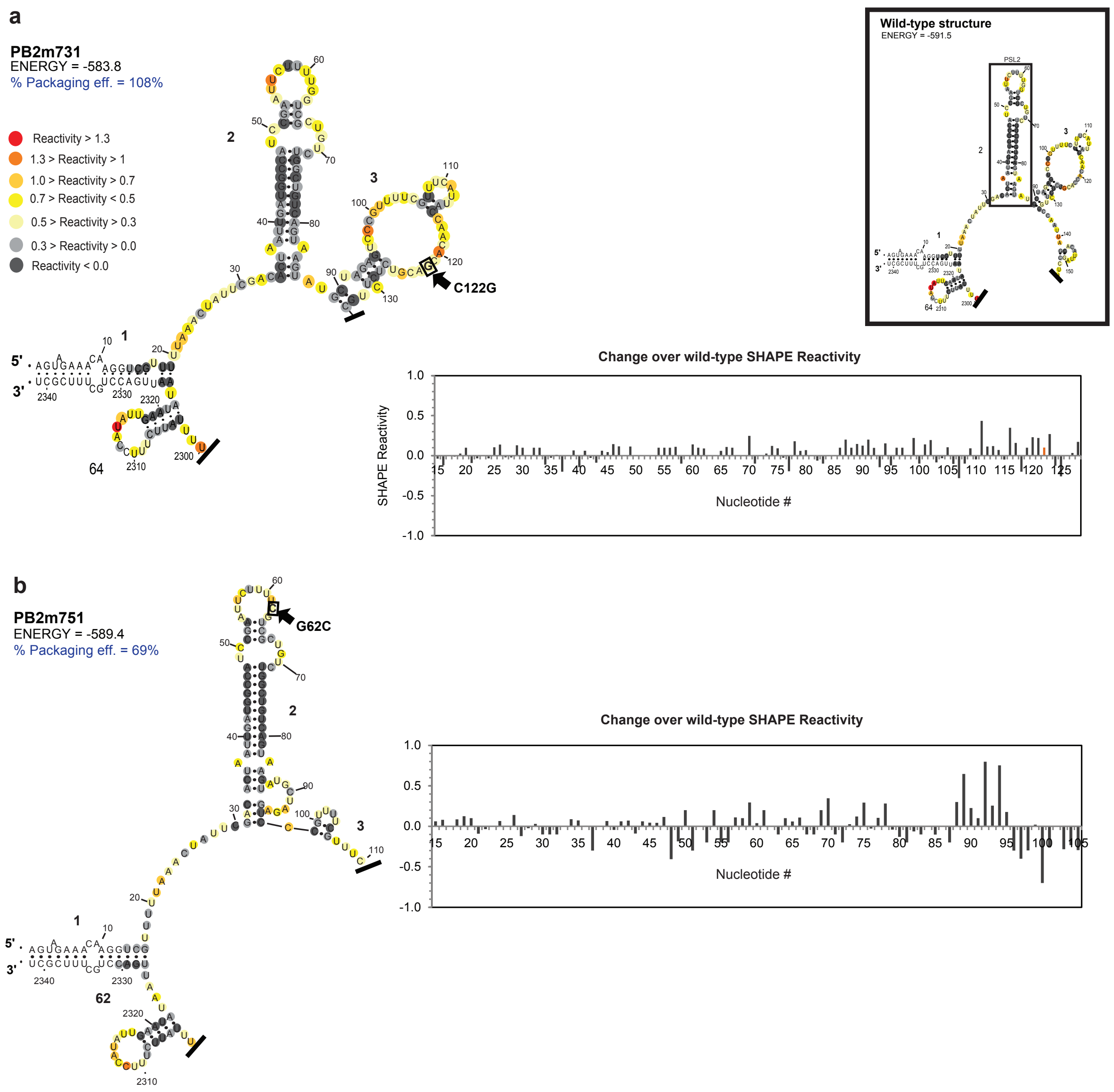


**Supplementary Figure 5. Effect of synonymous mutation on PSL2 structure.** Left: Predicted RNA secondary structure of PB2 packaging mutants determined by SHAPE analysis on full-length (-)-sense PB2 vRNA from PR8 strain. For reference, the wild-type structure is shown in upper right-hand corner box. Right: SHAPE reactivity graph shown as change of mutant reactivity over wild-type. Energy values and percent packaging efficiencies indicated below figure headings. Orange bars indicate site of mutation. Mutants: (**a**) m731. (**b**) m751. Percent packaging efficiencies of PB2 incorporation for each of the previously described mutants are highlighted in blue.

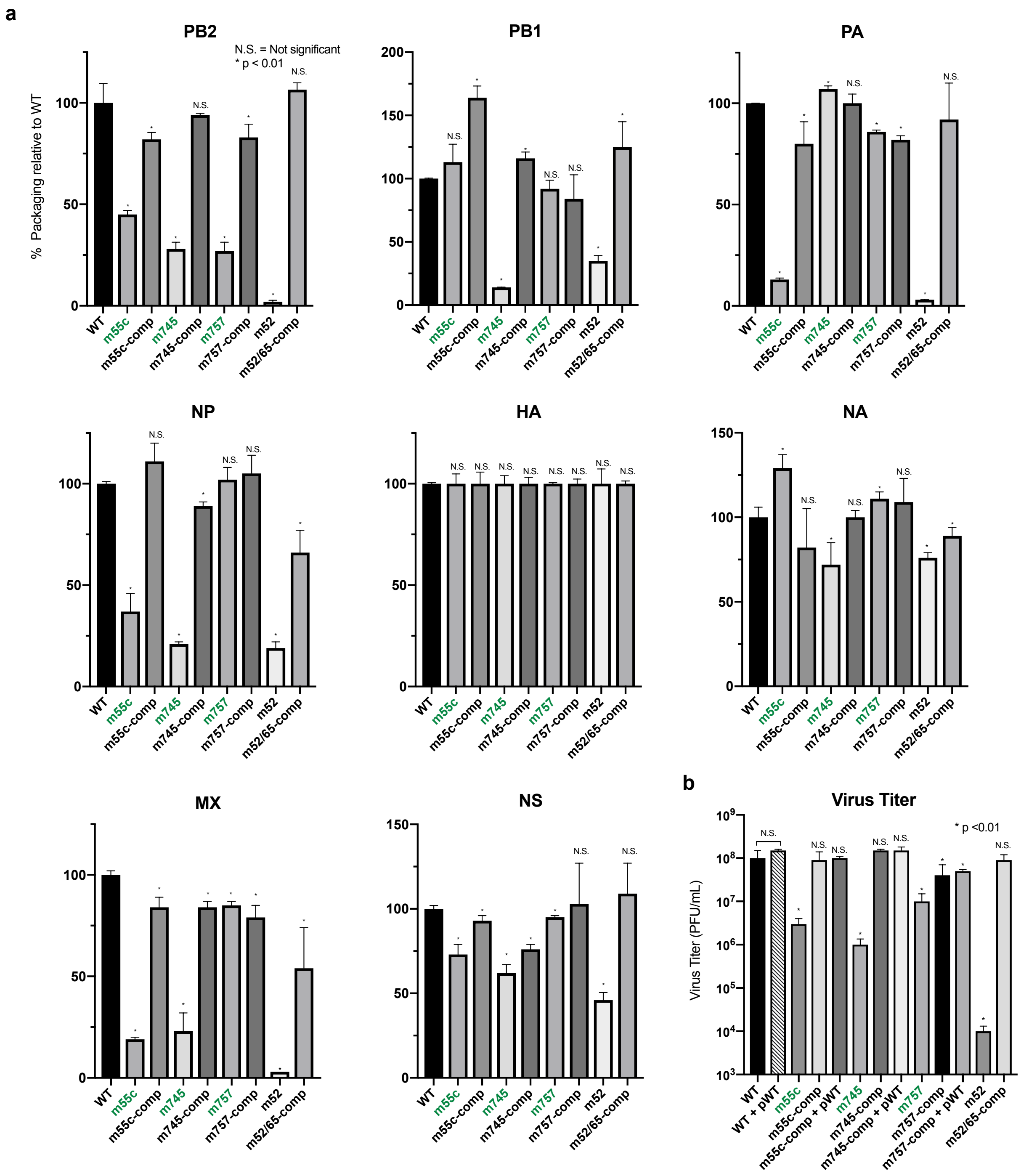


**Supplementary Figure 6. Effect of defective and compensatory mutations on IAV segment packaging and virus titer.** (**a**) Packaging efficiencies of defective and compensatory PB2 mutant viruses assayed for all eight IAV segments by qPCR. Previously described mutants are bolded in green. For compensatory mutations where a non-synonymous change was required, a wild-type PB2 protein expression plasmid was co-transfected during virus rescue. pWT = expression plasmid encoding for wild-type PR8 PB2 protein. Rescued viruses were amplified in eggs. Virus titers and packaging efficiency were measured from the harvested and clarified egg supernatant. Values given as percentage of segment vRNA packaging normalized to HA packaging efficiency relative to wild-type parental PR8 virus. Results from two independent experiments, assays performed in triplicate (n=6). (**b**) Virus titer determined by plaque assay. Results in PFU/mL, plaque assays performed in triplicate. Error bars represent ± s.d. Statistical significance determined by ordinary one-way ANOVA using Dunnett's multiple comparisons against wild-type, p values as indicated.

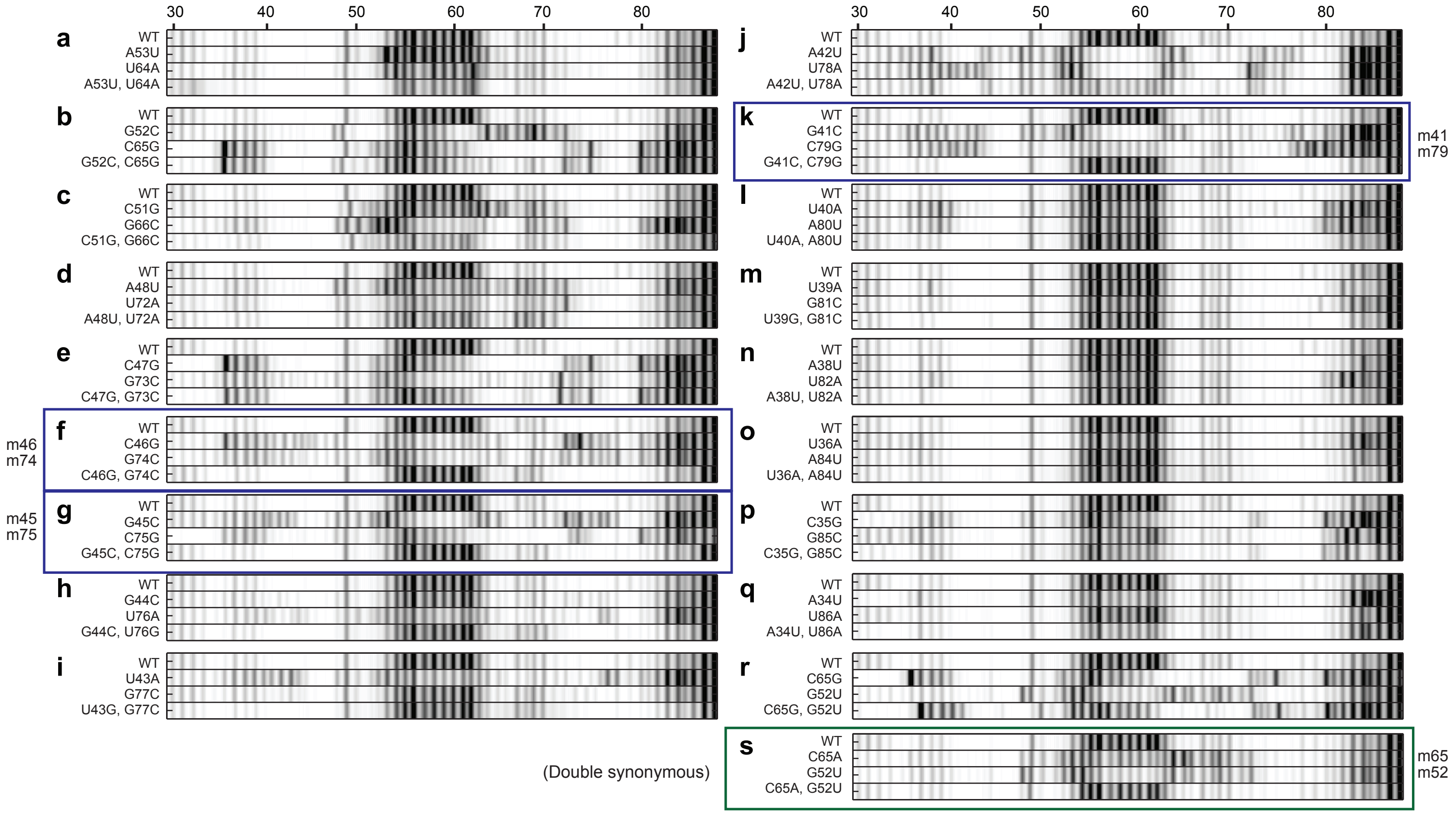


**Supplementary Figure 7. Mutate-Map-Rescue ($M^2R$) analysis.** Mutation/Rescue results validate PSL2 RNA secondary structure. Electropherograms of SHAPE analysis with compensatory double mutations to test base-pairings from 1D-data-guided models and to identify successful PSL2-defective and compensatory mutant pairs. Chemical accessibilities, plotted in grayscale (black = highest SHAPE reactivity), across 59 single mutations at single-nucleotide resolutions of PSL2 element from PR8 strain segment PB2. For each tested pairing, a "quartet" of wild-type, single mutant 1, single mutant 2, and compensatory double mutant are grouped for comparison. (**a-r**) Non-synonymous mutate-and-rescue pairs. All non-boxed electropherograms are pairings for which disruption and/or rescue were not observed. Blue boxes indicate successful defective and rescue mutations. (**s**) Double synonymous mutate-and-rescue pair (Green box).

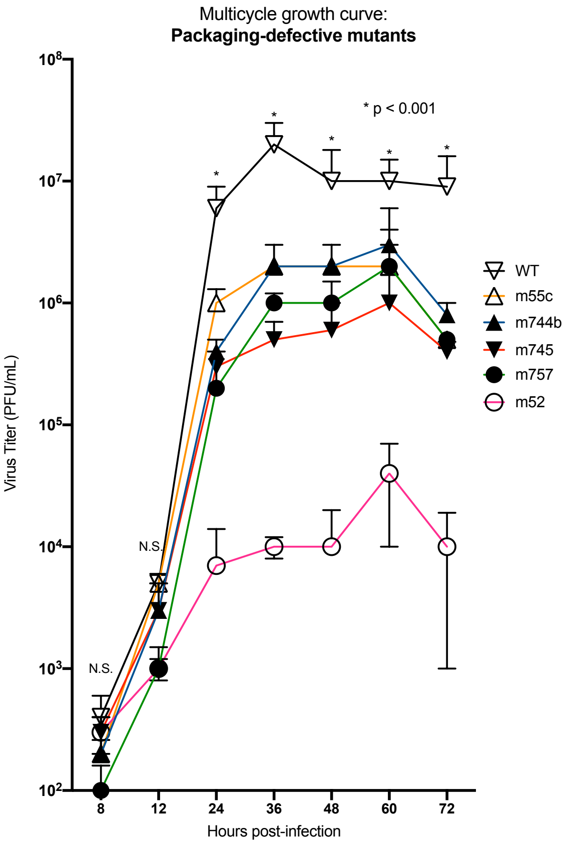


**Multicycle growth curve.** MDCK cells were infected at an MOI of 0.01 with either wild-type PR8 virus or packaging-defective mutant viruses. At the time points indicated, supernatant from biological replicates were collected and plaque assay titered in triplicate (n=6). Error bars represent ± s.d. Statistics performed by multiple comparisons via 2way ANOVA; p value determined between WT control and mutants,* p < 0.001.

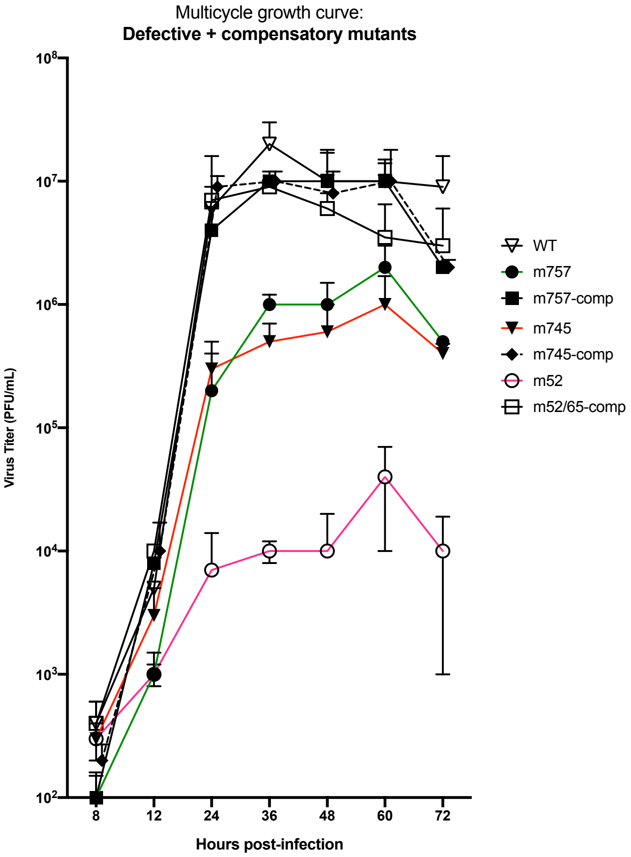


**Multicycle growth curve.** MDCK cells were infected at an MOI of 0.01 with either wild-type PR8 virus or packaging-defective or compensatory mutant viruses. At the time points indicated, supernatant from biological replicates were collected and plaque assay titered in triplicate (n=6). Error bars represent ± s.d.

**Supplementary Figure 8. Multi-cycle growth curve of packaging-defective viruses.** MDCK cells were infected at an MOI of 0.01 with either wild-type PR8 virus or packaging mutant viruses. At the time points indicated, supernatant from biological replicates were collected and plaque assay titered in triplicate. Error bars represent ± s.d. Statistical analysis performed by multiple comparisons via 2way ANOVA; p value determined between WT control and mutants, p * < 0.001.

| | | |
|---|---|---|
| (Syn.) **m52-F (C2290A):** | CCA AAA GAA TTA GGA TGG CCA TCA ATT AGT GTC G | Double synonymous mutant pair |
| (Syn.) **m52-RC (C2290A):** | CGA CAC TAA TTG ATG GCC ATC CTA ATT CTT TTG G | |
| (Syn.) **m65-F (G2277T):** | GAC AGC CAG ACA GCT ACC AAA AGA ATT CG | |
| (Syn.) **m65-RC (G2277T):** | CGA ATT CTT TTG GTA GCT GTC TGG CTG TC | |

**Supplementary Figure 9. Design of primer sequences for 2-Dimensional Mutate-Map-Rescue ($M^2R$) mutants.** Primer sequences used for QuickChange mutational cloning of $M^2R$ mutants into pDZ plasmids. Sequences are in (+)-sense orientation. Left field denotes synonymous (Syn.) or non-synonymous (Non-syn.) change. Highlighted nucleotides = mutation site. Boxed mutant primer set indicates double synonymous mutant partners, m52 and m65.

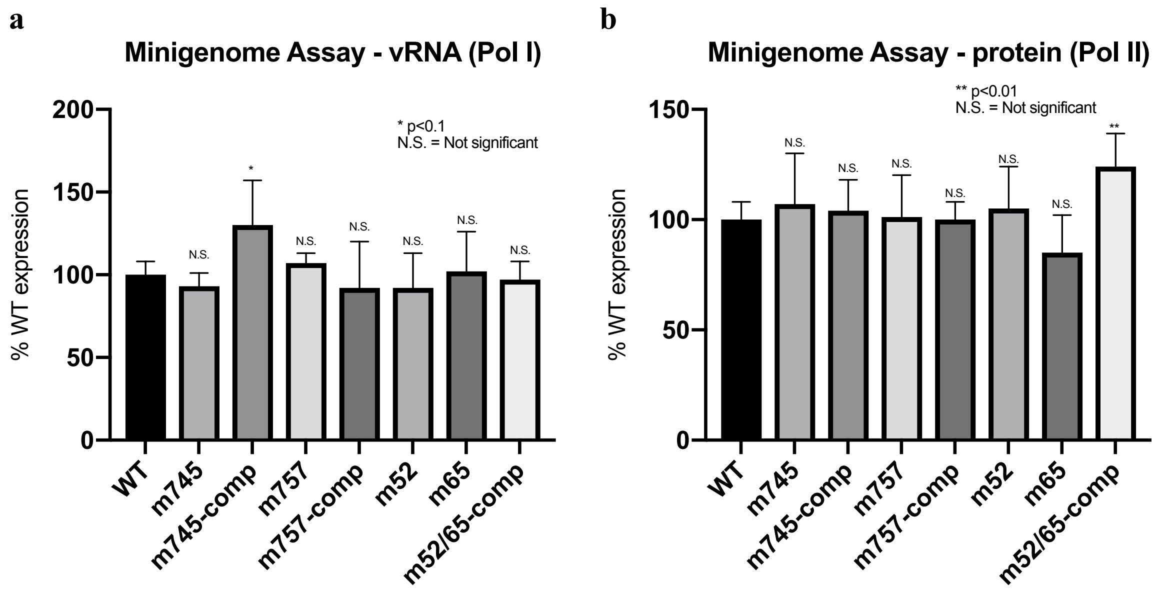


**Supplementary Figure 10. Replicon assay of defective and compensatory PB2 packaging mutants.** (**a**) To test the ability of mutant PB2 vRNA to be replicated, wild-type protein expression plasmids (Pol II-directing) for the viral polymerase genes comprising PB2, PB1, PA, and NP segments were cotransfected into 293T cells with either a wild-type or mutant PB2 vRNA (Pol I-directing) plasmid, in addition to a wild-type HA vRNA-directing Pol I plasmid that served as transfection control. (**b**) To test the effect of packaging mutations on the ability of the mutant PB2 protein to function as part of the polymerase complex, wild-type protein expression plasmids for PB1, PA, and NP were cotransfected with mutant PB2 protein expression plasmids (Pol II-directing) and wild-type PB2 vRNA plasmid (Pol I). Wild-type HA vRNA plasmids were used as transfection controls. Twenty-four hours post-transfection, RNA was isolated and extracted by TriZol. Results were readout by qPCR, normalized to HA as the transfection control, and are shown relative to wild-type expression. Assays performed in triplicate. Error bars represent ± s.d.

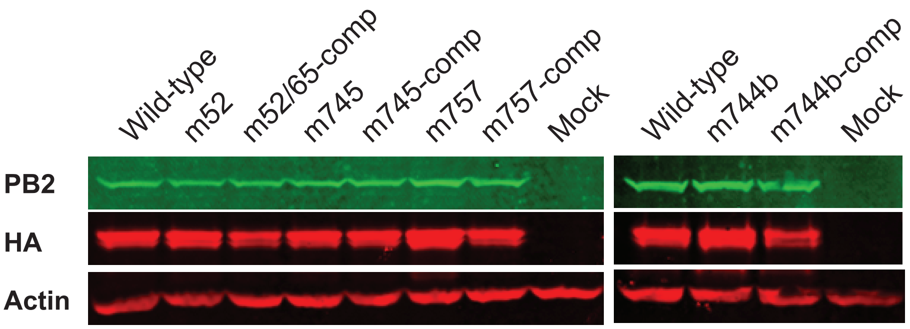


**Supplementary Figure 11. Western blot of mutant PB2 protein expression in virus infected cells.** MDCK cells were infected with either wild-type or packaging mutant PR8 viruses at high MOI (6) to achieve a single-cycle infection. Six hours post-infection cells were washed with ice cold PBS twice and harvested in Laemmli sample buffer followed by DNase treatment to digest genomic DNA. The samples were boiled for 5 min and equal amounts of cell lysate were subjected to SDS-PAGE and western blotting. The membranes were probed with Rabbit anti-PB2 (Invitrogen) followed by anti Rabbit-IRDye800 (Rockland), mouse anti-HA (Sinobiological) followed by anti-mouse AlexaFluor-680 (Invitrogen), and mouse anti-Actin (Sigma) followed by anti-Mouse AlexaFluor680. The blot was probed separately for each antibody and scanned in an Odyssey CLx scanner.

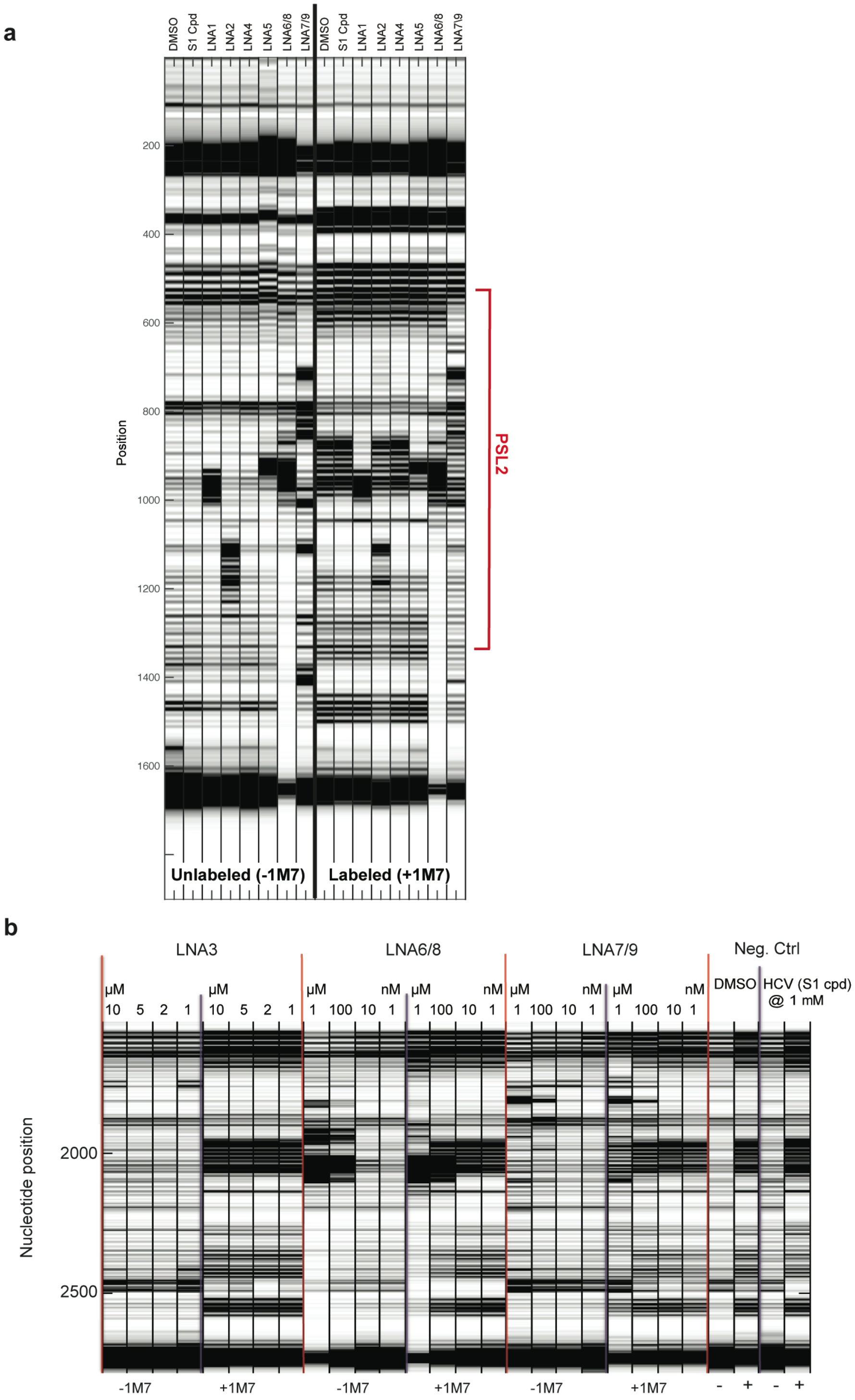


**Supplementary Figure 12. SHAPE analysis on LNA-RNA binding.** (**a**) Electrophoretic profile of SHAPE analysis performed on LNAs 1,2, 4, 5, 6/8, and 7/9 (100 nM) in the presence of PR8 PB2 vRNA. S1, a small molecule that interacts with the Hepatitis C virus IRES RNA was added as a control for RNA binding. Columns on the left, unlabeled reaction without SHAPE reagent, (-1M7). Right: with labeling reagent (+ 1M7). (**b**) Electrophoretic profile of SHAPE performed on LNA-vRNA combination in titrating concentrations of LNA. For each LNA, the left set of columns is without labeling reagent. Positions of PSL2 indicated by red brackets.

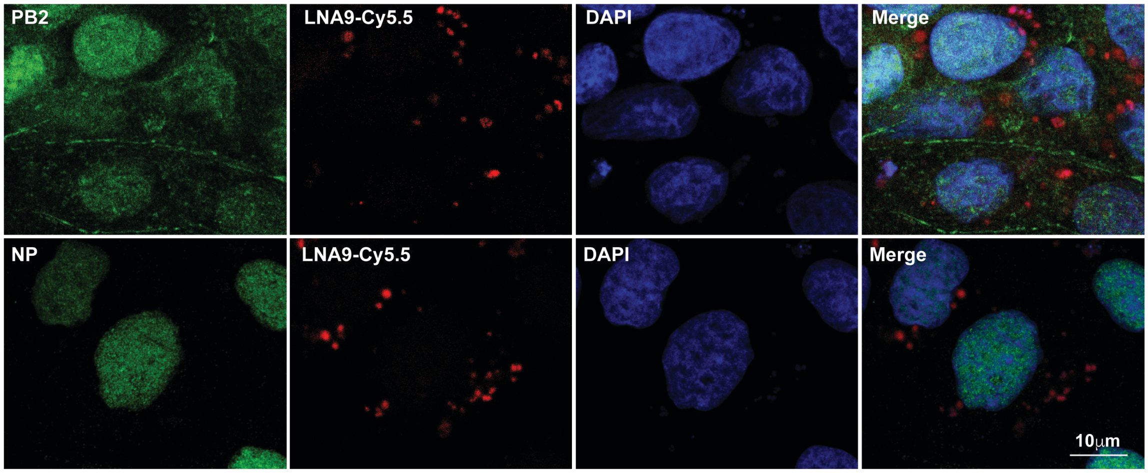


**Supplementary Figure 13**. **Immunofluorescence analysis of LNA9 and vRNP in LNA treated infected cells.** MDCK cells were transfected with Cy5.5-labeled LNA9 four hours prior to infection with PR8 virus (MOI 0.01). 15 hours post infection, cells were rinsed with phosphate-buffered saline twice, fixed in 4% fresh paraformaldehyde and permeabilized with 0.2% Triton X-100. After blocking in Tris buffered saline (TBS) containing 10% goat serum, cells were stained using rabbit anti-PB2 (Invitrogen) for 45 min. After several washing steps, cells were stained with Alexa 488–conjugated goat anti-rabbit IgG or anti-mouse IgG (Invitrogen) for an additional 45 min. Slides were mounted with prolong gold anti-fade reagent with DAPI (Invitrogen), prior to examination under Zeiss LSM 510 confocal microscope (Carl Zeiss MicroImaging, Thornwood, NY).

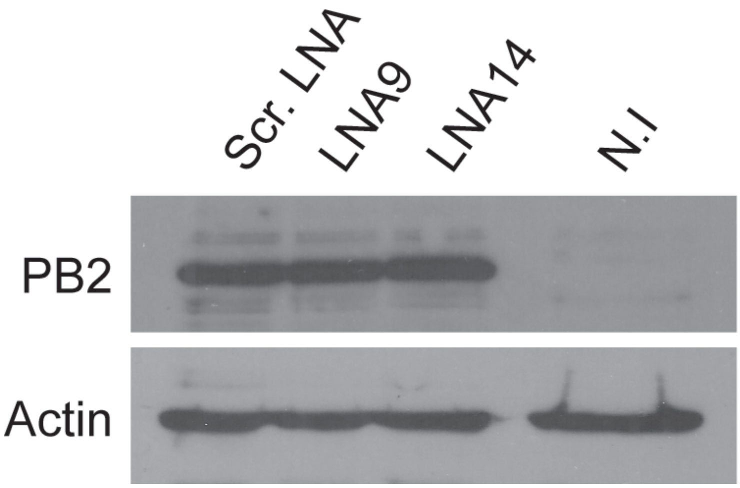


**Supplementary Figure 14**. **Western blot analysis of LNA9 and scrambled LNA treated infected cells.** MDCK cells were transfected with 1 μM Scrambled LNA or PSL2 targeting LNA9. Twenty-four hours post transfection, the cells were infected with PR8 virus at an MOI of 0.1. The cells were harvested 15 hours post-infection. Equal amounts of total cell lysate were resolved on a 10 % SDS-PAGE gel followed by transfer to PVDF membranes. The membrane was blocked with 5 % non-fat dry milk in TBST, incubated with rabbit anti-PB2 or mouse anti-actin primary antibodies, followed by the appropriate secondary antibodies conjugated to HRP. A chemiluminescent detection of the proteins was performed and films were developed using a Kodak film developer. N.I. = non-infected cells.

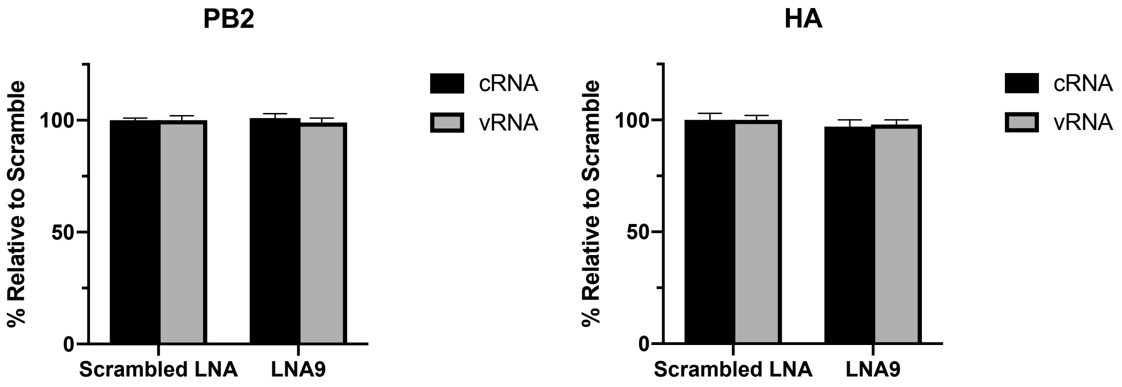


**Supplementary Figure 15**. **qPCR analysis of vRNA and cRNA in LNA9 and scrambled LNA treated infected cells.** MDCK cells were pretreated with 100 nM of LNA9 or Scrambled LNA. Eight hours post-transfection, the cells were infected with PR8 virus at an MOI of 1. Eight hours post-infection, cells were harvested and total RNA extracted. Influenza vRNA and cRNA for PB2 and HA genes were measured by strand-specific RT-qPCR. RNA levels normalized to scrambled LNA treated cells are presented. Error bars represent ± s.d.

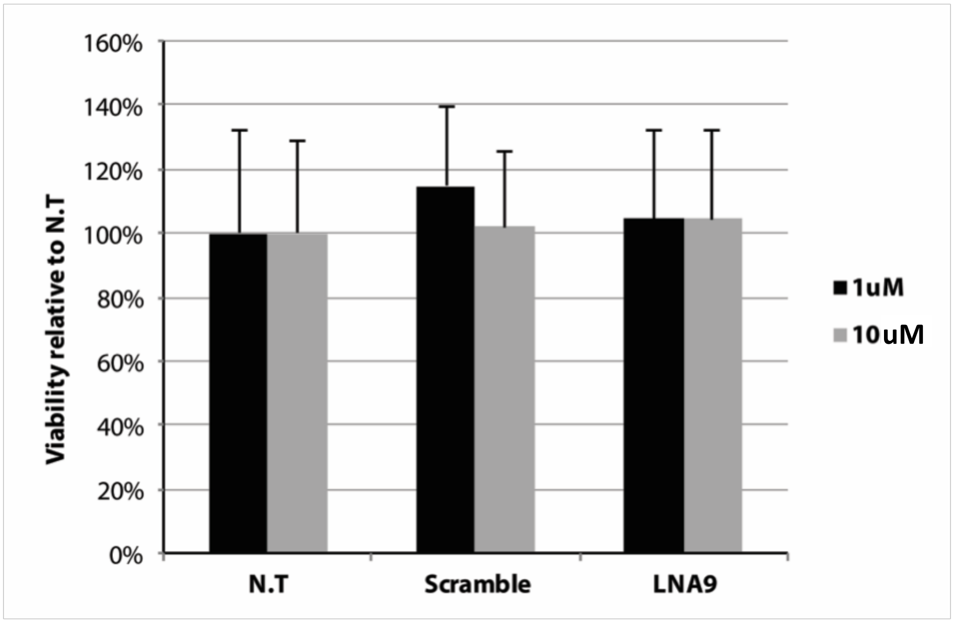


**Supplementary Figure 16**. **LNA9 treatment of cells has no cellular cytotoxicity.** MDCK cells were treated with media control vs. 1 or 10 micromolar scrambled LNA or LNA9 followed by treatment with PrestoBlue to assess for cellular toxicity. Results expressed as a percent of non-treated control.

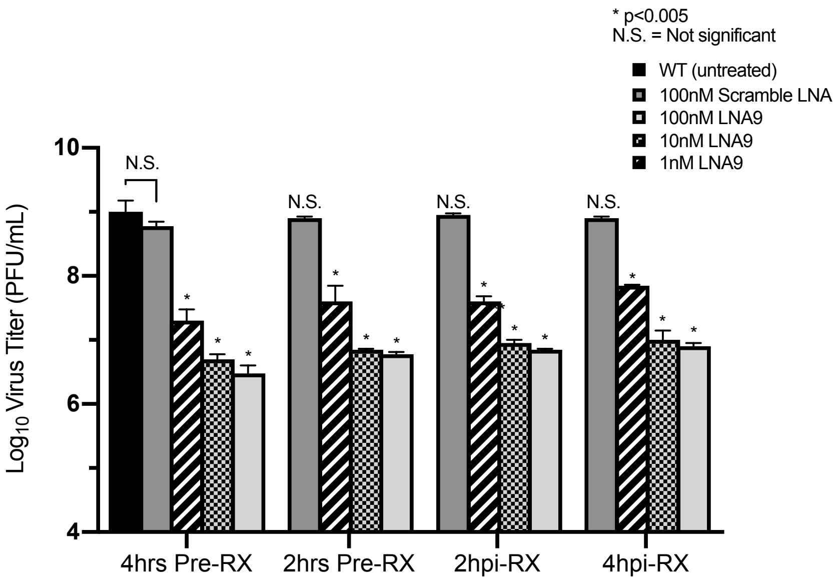


**Supplementary Figure 17**. **LNA9 treatment of cells infected at high MOI**. Time course of pre-treatment (Pre-RX) versus post-infection treatment with scrambled LNA and non-treated control vs. LNA9 at titrating concentrations (100 nM, 10 nM, 1 nM). Pretreatment: MDCK cells were treated with LNA9 either 2 or 4 hours prior to infection with 0.1 MOI of PR8 virus. Post-infection treatment: MDCK cells were infected with 0.1 MOI of PR8 virus and treated with scrambled LNA and the indicated concentrations of LNA9 at either 2 or 4 hours post-infection. Supernatant was collected 48 hpi, and viral titers determined by plaque assay. Error bars represent ± s.d. *=p <0.05 relative to scramble control.

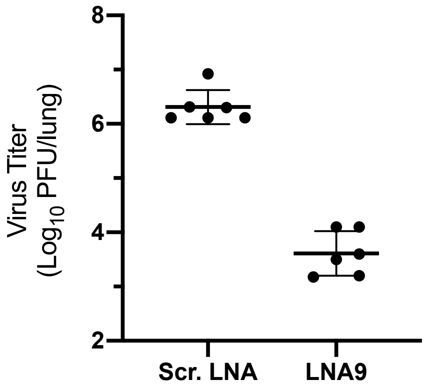


**Supplementary Figure 18**. **Lung virus titers of LNA-treated mice post-infection.** BALB/c female mice were treated with 20 µg of LNA9 or Scrambled LNA (Scr. LNA) twenty-four hours prior to infection with 800 PFU of PR8 virus (Day 0). At Day 3 post-infection, each group was sacrificed, and whole lungs were harvested, weighed and homogenized in PBS proportional to total lung weight. Viral titers of the homogenized lung supernatant were determined by plaque assay then normalized to lung weight. Three mice/group, plaque assays performed in duplicate (N=6). Error bars represent ± s.d.

a

#### Mutant Nomenclature and Mutation Sites

| | Mutant Name | Mutation Site(s) [(-)-sense orientation] | SYN or Non-SYN | Notes |
|---|---|---|---|---|
| Previously-described Mutants | **m55c** | CU35,36UC | SYN | |
| | **m757** | G44C | SYN | |
| | **m55a** | CAU50,53,56 | SYN | |
| | **m751** | G62C | SYN | |
| | **m748** | C71U | SYN | |
| | **m745** | A80U | SYN | |
| | **m744b** | AG83,85UA | SYN | |
| | **m737** | U104 | SYN | |
| | **m731** | C122G | SYN | |
| Novel PSL2-informed Mutants | **m83,84** | AG83,84GA | Non-SYN | |
| | **m55c-comp** | CU35,36UC / AG83,84GA | Non-SYN | Dbl Mutant |
| | **m76** | U76G | Non-SYN | |
| | **m757-comp** | G44C / U76G | Non-SYN | Dbl Mutant |
| | **m40** | U40A | Non-SYN | |
| | **m745-comp** | A80U / U40A | Non-SYN | Dbl Mutant |
| | **m74-1** | G74A | SYN | |
| | **m74-2** | UCG74AGC | SYN | |
| | **m68** | U68A | SYN | |
| | **m56** | U56G | SYN | |
| | **m41** | G41C | Non-SYN | |
| | **m79** | C79G | Non-SYN | |
| | **m41/79-comp** | GC41 / 79CG | Non-SYN | Dbl Mutant |
| | **m45** | G45C | Non-SYN | |
| | **m75** | C75G | Non-SYN | |
| | **m45/75-comp** | GC45 / 75CG | Non-SYN | Dbl Mutant |
| | **m46** | C46G | Non-SYN | |
| | **m74** | G74C | Non-SYN | |
| | **m46/74-comp** | CG46 / 74GC | Non-SYN | Dbl Mutant |
| | **m52** | G52U | SYN | |
| | **m65** | C65A | SYN | |
| | **m52/65-comp** | GC52 / 65UA | SYN | Dbl Mutant |

**SYN = Synonymous mutation, Non-SYN = non-synonymous mutation**

b

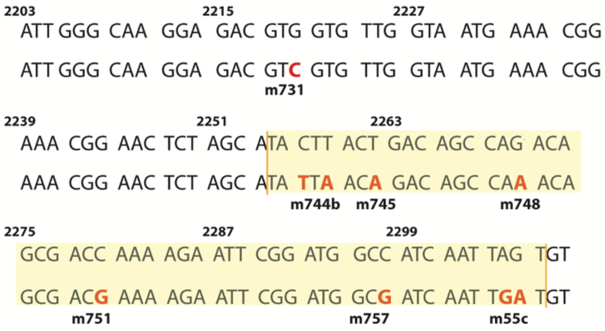


**Supplementary Table 1. PB2 Packaging mutant nomenclature and corresponding sites of mutation**.

(**A**) Mutation nomenclature chart indicating names and site(s) of mutation from: 1) Previously published synonymous mutations implicated in PB2 packaging (shown in blue) (*13, 16*); and 2) PSL2 structure designed single mutants (shown in black) and double-compensatory mutants (shown in red). Numbering and nomenclature of introduced mutations are based on the genomic, (-)-sense vRNA. Instances where mutation results in change in protein coding are indicated by synonymous (SYN) or non-synonymous (Non-SYN) fields. (**B**) Previously published synonymous mutations implicated in PB2 packaging. Upper line is the parental PR8 vRNA sequence ((+)-sense orientation), and the mutated single nucleotides are bolded in red on the line below. Numbering and nomenclature of introduced mutations are based on the reports by Marsh et al. (*13*) and Gog et al. (*16*). Yellow highlighted region indicates sequence containing PSL2 structure.

### Clinical Score Calculation

| Score | Coat | Respiration | Mobility | Posture | Weight |
|---|---|---|---|---|---|
| **1** | Slightly scruffy / in some parts | Rapid | Slower / Reduced | Slightly hunched | 20% |
| **2** | Rough fur / on whole body | Difficult / labored breathing (Dyspnea) | Slow / None | Very hunched / non responsive | 25% |
| **3-4** | Any combination of the above | | | | |
| **5** | Any combination of the above **OR** if weight loss >30% | | | | |

Clinical score: sum of the value for these conditions results with a clinical score

Humane euthanization: When a clinical score is 5 and up OR 30% body weight loss

**Supplementary Table 2. Description of clinical scoring in diseased mice.**
